# Sequential transcriptional gates in the thalamo-cortical circuit coordinate memory stabilization

**DOI:** 10.1101/2025.05.28.656415

**Authors:** Andrea Terceros, Celine Chen, Yujin Harada, Tim Eilers, Millennium Gebremedhin, Richard Koche, Pierre-Jacques Hamard, Roshan Sharma, Priya Rajasethupathy

**Author notes:** These authors contributed equally (Andrea Terceros, Celine Chen).

## Abstract

The molecular mechanisms that enable memories to persist over long time-scales from days to weeks and months are still poorly understood. To develop insights we created a behavioral task where, by varying the frequency of learned associations, mice formed multiple memories but only consolidated some, while forgetting others, over the span of weeks. We then monitored circuit-specific molecular programs that diverge between consolidated and forgotten memories. We identified multiple distinct waves of transcription, i.e., cellular macrostates, specifically in the thalamo-cortical circuit, that defined memory persistence. Notably, a small set of transcriptional regulators orchestrated broad molecular programs that enabled entry into these macrostates. Targeted CRISPR-knockout studies revealed that while these transcriptional regulators had no effects on memory formation, they had prominent, causal, and strikingly time-dependent roles in memory stabilization. In particular, the calmodulin-dependent transcription factor Camta1 was required for initial memory maintenance over days, while Tcf4 and the histone methyl-transferase Ash1l were required later to maintain memory over weeks. These results identify a critical Camta1-Tcf4-Ash1l thalamo-cortical transcriptional cascade required for memory stabilization, and puts forth a model where the sequential, multi-step, recruitment of circuit-specific transcriptional programs enable memory maintenance over progressively longer time-scales.

---

Memories are maintained across vastly different time-scales, from hours, to days, to months and years. The molecular mechanisms by which memories are stabilized over progressively longer time-scales are still poorly understood. A key insight came from early observations that transcriptional blockers, while leaving short-term memories intact, prevented the formation of longer-term memories, from honeybees to goldfish to mice^1–4^. These and other studies thus revealed that the synthesis of new proteins is required to prolong hour-long memories^5^ to days-long memories^6–11^. One transcription factor, CREB, has been implicated in this process, where its suppression prevents the formation of long-term memories, whereas its activation can potentiate the conversion of a transient into an overnight memory^12–17^. Thus, the role of transcription in sensing transient signals and activating genes that can prolong functional and structural changes at synapses provides a framework for extending memory from hours to days (termed synaptic consolidation) ^18–20^. However, the molecular programs recruited in extended brain circuits that enable memories to persist on longer time-scales, over weeks, months, or even a lifetime (termed systems consolidation), are yet unknown.

Prior work has detailed that memories can be stabilized through a process of consolidation and reconsolidation^21–23^ suggesting that additional transcriptional programs may be recruited to extend memories over longer time-scales. Furthermore, epigenetic factors have important roles in maintaining cellular memory during development for weeks and months, for instance as cell lineages are specified and maintained, which are processes that may be co-opted in adult neural circuits for prolonging memories^24–26^. Finally, local protein synthesis^27,28^ and long-lived enzymatic and structural changes^29^ may also work in concert to extend memory persistence.

To bridge these models and gain insights into the longer time-scale maintenance of memories, we developed a behavioral task where some memories are consolidated while others are forgotten over the span of weeks. We then developed approaches to study evolving cellular transcriptomes that are unique to consolidated memories, followed by loss-of-function gene manipulations to assess their impact on memory maintenance. In doing so, we identified discrete waves of transcription in the thalamo-cortical circuit, governed by specific non-canonical memory-related transcriptional regulators, that are required for the progressive stabilization of memories spanning days to weeks.

## A behavioral task to monitor memory persistence

To identify molecular programs uniquely associated with memory persistence, we began by developing a behavioral task where mice form multiple memories but only consolidate some while forgetting others over the span of weeks. Since repetitive exposure during learning affects memory persistence, we trained mice to learn context-outcome associations that were presented at varying frequencies. Thus, mice were presented with two reward associated contexts, one at high repetition (HR, ∼50% of trials) and the other at low repetition (LR, ∼22% of trials), with the expectation that mice will form both memories but only maintain memory of the HR context (i.e., by licking in anticipation for the associated reward) but not of the LR context over weeks. To avoid continuous non-associative licking, we randomly interleaved the two reward contexts with an aversive context (A, ∼28%) for a total of ∼50 trials per session, over a ∼7 day learning period (Fig. 1a, top). To tightly control the frequency of context presentations, and record behavioral outcomes at high temporal precision, while maximizing the number of trials collected, we implemented this behavior as a head-fixed virtual-reality based task.

**Fig. 1:**
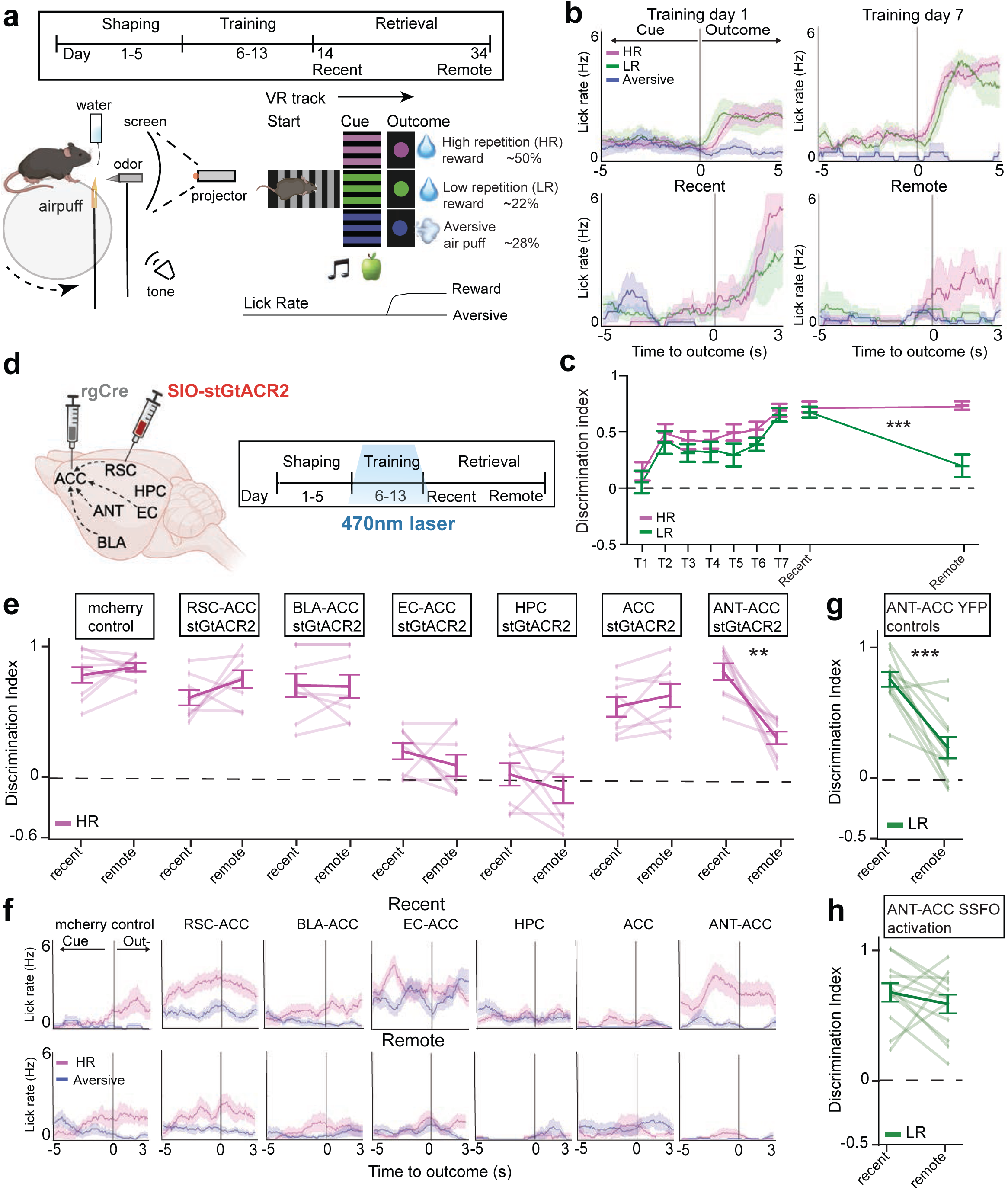
A behavioral task to monitor memory persistence requires the HPC, ANT and ACC. **a**, Top: timeline of behavioral task from shaping (day 1), through training (days 6–13), and retrieval days (days 14 and 34). Bottom left: schematic of virtual reality experimental setup. Bottom right: virtual reality linear track with start, cue, and outcome zones, and example lick rate tracked. **b**, Representative lick traces from one mouse per cohort showing trial averages on first and last days of training (top) and recent and remote retrieval; no reinforcement given, bottom, *n* = 40-50 trials per session. Data are mean (solid line) ± SEM (shaded area). **c**, Quantification of discrimination indices for learning and retrieval in HR and LR, *n* = 24 mice, data are mean ± SEM, \*\*\**P* = 0.000062 between recent and remote for LR, paired t-test with Bonferroni-Dunn correction. **d**, Top: injection strategy to target projections to ACC from ANT, RSC, EC and BLA in SIO-stGtACR2 opsin cohorts. Local stGtACR2 injected in HPC and ACC. Light delivered during cue periods of training sessions only, and memory was tested on recent and remote time points. **e**, Quantification of discrimination indices between HR and aversive lick rates, *n* = 7-9 mice per cohort, individual data points shown, \*\**P* = 0.0057 for ANT-ACC SIO-stGtACR2, paired t-test with Bonferroni-Dunn correction. **f**, Representative raw lick traces from one mouse for each of the cohorts summarized in d, *n* = 30-40 trials, data are mean (solid line) ± SEM (shaded area). **g**, Quantification of discrimination between low reward (LR) and aversive for YFP (no opsin control, n = 10) on mid and remote retrieval, \*\*\**P* = 0.00048 paired t-test with Bonferroni-Dunn correction. **h**, Same as in (g) but for ANT-ACC SSFO excitation, n = 12.

In brief, mice navigated on an axially fixed track ball in a virtual-reality environment composed of a corridor with three distinct zones: start, cue, and outcome zone. Trials were initiated in the start zone. Then, mice entered one of three cue zones (each being a unique multimodal context consisting of auditory, visual, and olfactory cues), two of which were paired to a water reward in the outcome zone (HR & LR), while the other was paired to an aversive air puff (A) (Fig. 1a, bottom). The cues for each context were designed to be multi-modal and spatial in nature to ensure hippocampal dependency during memory formation (Methods). By the end of training, mice reliably learned the context associations by exhibiting anticipatory licking in contexts that predicted reward while suppressing licking in the context that predicted an aversive air puff (Fig. 1b, top). During the retrieval phase, mice were presented with probe trials at a recent (Day 1) and remote (Day 21) time, where reward and aversive outcomes were omitted, and lick rates in the outcome zone provided a measure of recall in the absence of reinforcement. Mice demonstrated successful recall of both the HR and LR contexts at recent time, as evidenced by significant and sustained differences in raw lick rates in outcome zones, as well as by their lick discrimination index (Figure 1b and 1c). Over time, they preferentially maintained memory of the HR context, while failing to recall the LR context by remote time (Figure 1c, *P*<0.001 between recent and remote for LR, paired t-test with Bonferroni-Dunn correction). These differential lick rates did not exist when cue-outcome pairs were shuffled (Extended data Fig. 1a), ensuring that mice did not display intrinsic lick preferences for one context over the other, but rather formed learned associations.

We next asked whether this task requires mice to use brain circuits known to be involved in memory formation and consolidation. While the formation of contextual memories initially requires the hippocampus, the subsequent days to weeks long consolidation process becomes increasingly dependent on cortical structures, and in particular the anterior cingulate region of the prefrontal cortex^30^. We therefore conducted loss of function experiments to test the hippocampal and cortical dependency of memory in this task. We expressed the inhibitory opsin stGtACR2 bilaterally in either the hippocampus (HPC, CA1 region) or anterior cingulate cortex (ACC, layers 2/3) and delivered light in the cue zones to silence these regions during recent or remote retrieval. As expected, inhibition of the HPC during retrieval led to near complete deficit in memory recall at recent time, whereas inhibition of ACC resulted in intact recent memory but a strong remote memory deficit (extended data fig. 1b, 1d). These results confirm that this behavioral task captures the temporal shift in hippocampal-to-cortical dependency during weeks-long memory consolidation.

We previously demonstrated that the anterior thalamus (ANT) is an important conduit that supports hippocampal-to-cortical memory consolidation^31^. We thus asked whether the ANT to ACC projection is required for memory consolidation in this new task, and we simultaneously took a broad survey to assess the relative contributions of other known ACC projecting pathways to memory consolidation. We expressed AAVretro-Cre in ACC and SIO-stGtACR2-mCherry (or Floxed-mCherry control) bilaterally in ANT, retrosplenial cortex (RSC), basolateral amygdala (BLA), or entorhinal cortex (EC) in separate cohorts of mice. We then performed projection specific silencing of each of these pathways during training and tested their effect on recent and remote recall. To assess the magnitude of these effects, we also included cohorts of mice expressing bilateral stGtACR2 in HPC and ACC for comparison. Overall, we observed that while some manipulations produced strong deficits in memory formation (HPC, EC>ACC,), others produced neither deficits in memory formation nor consolidation (RSC>ACC, BLA>ACC, ACC) (Fig. 1e, representative raw lick traces in Fig. 1f, extended data Fig. 1c). Notably, only the ANT>ACC manipulation produced intact learning and recent memory but an isolated deficit in memory consolidation. This effect size was particularly striking with fully intact recent recall (DI ∼0.7), comparable to controls, but near complete deficit in recall by remote time (DI ∼0.1) (Fig. 1e, *P* = 0.004 for recent vs remote, paired t-test with Bonferroni-Dunn correction). While we do not rule out the contribution of other circuits (they may function at different times or have mixed roles during learning/memory thus precluding an isolated consolidation deficit), these results support a prominent role for the HPC to ANT to ACC pathway in memory consolidation. Additionally, we tested whether activation of the ANT-ACC circuit during training would be sufficient to improve recall of LR at remote time (which we previously found to be true^31^ but in a different task where LR was defined by low reward *value* rather than low *repetition*). We expressed the stabilized step-function opsin (SSFO) specifically in the ANT-to-ACC circuit and delivered light only during training. We found that while this manipulation had no effect on learning, it had a striking ability to enhance LR memory at remote time, which would otherwise have been forgotten (Fig. 1g, 1h, *P* < 0.001 for recent vs remote, paired t-test with Bonferroni-Dunn correction). We thus established a behavioral task, and its neural circuit dependencies, that would allow us to study the diverging molecular programs that enable some memories to persist while others are forgotten.

## Distinct time-dependent transcriptional programs are activated in the thalamus and cortex during memory maintenance

While molecular programs in the hippocampus that stabilize overnight memories have been well studied, relatively little is understood about molecular programs that extend memories from days into weeks^32–34^. Given the requirement of the ANT-ACC circuit in supporting days-to-weeks-long memories, we next studied how molecular programs in this circuit diverge between HR (eventually consolidated) and LR (eventually forgotten) over time. We reasoned that the process of memory consolidation may drive some neurons into distinct cellular states, of varying durations, uniquely in the HR condition. To test this, we performed single-cell RNA-sequencing (scRNA-seq) in the ANT-ACC circuit, at repeated time points through the course of the behavior, to capture the evolving molecular processes at cellular resolution.

We trained a cohort of 48 mice on the above-described behavioral task, then split the cohorts during retrieval such that each cohort ended in a terminal training (T) or retrieval (R) time point. Furthermore, at each retrieval time, the cohorts were further split into mice that only recalled HR versus mice that only recalled LR. In total this allowed at least n=3 biological replicates from each brain region (ANT, ACC), per time point (early-Training, late-Training, recent-retrieval, mid-retrieval and remote-retrieval) and per context (HR, LR) for single-cell dissociation and library preparation (Fig. 2a). We selected these time intervals based on prior work, which successfully mapped cellular transcriptomic trajectories using one week intervals to capture the developmental change of cell lineages^35,36^. To mitigate batch effects, we tracked sample identity with multiplexed barcodes and pooled all cells together for joint sequencing. As before, mice learned the HR and LR context associations equally well and performed similarly at recent retrieval, but the HR and LR cohorts exhibited strong divergence in behavioral recall by remote retrieval (extended data fig. 2a-2c). In total, we sequenced 76,566 cells from the ANT and 145,327 cells from the ACC. To improve statistical power in detecting cell types and phenotypic states, we merged all cells from all conditions and time points, and after standard preprocessing, we clustered all cells using the Phenograph algorithm^37^ and visualized the data in 2D using UMAP (extended data Fig. 2e, 2g). The resulting clusters were annotated into major cell types based on the average expression of canonical marker genes collected from published datasets^38,39^ (Fig. 2b, table S1). We focused all downstream analysis on the ANT and ACC neuronal clusters. As before, we were able to visualize the distribution of time points and biological replicates (extended data Fig. 2h, 2i, 2l, 2m). Sub-clustering of the ANT and ACC neurons (Fig. 2c, 2e) demarcated excitatory from inhibitory classes (extended data Fig. 2j, 2k, 2n, 2p), as well as anterior vs posterior nuclei in the ANT (Fig. 2d), and the distinct cortical layers in the ACC (Fig. 2f and extended data fig. 2o, 2q), based on the average expression of marker genes (table S2). Importantly, we ruled out batch effects by confirming similar representation of neuronal classes per condition and time point regardless of day of sample collection (extended data fig. 2k bottom, 2q bottom).

**Fig. 2:**
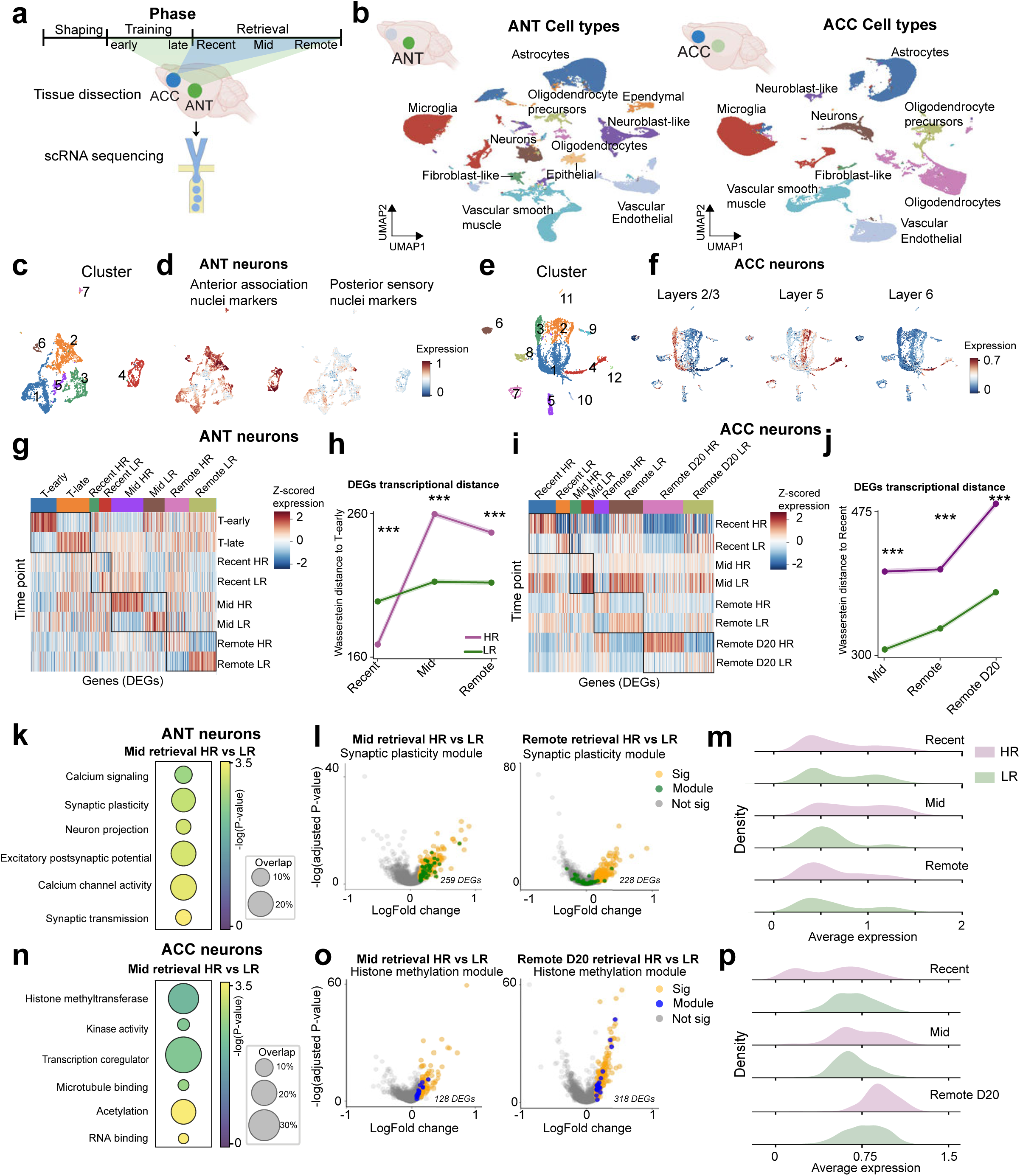
Distinct transcriptional programs are activated in the ANT and ACC during memory stabilization. **a**, Schematic representation of the scRNA sequencing workflow. **b**, Left: UMAP visualization of all cell types collected from ANT (*n* = 176566 cells from *n* = 23 mice), right: UMAP of all cell types collected from ACC (*n* = 145327 cells from *n* = 24 mice), clustered based on expression of canonical markers. **c, e**, UMAP sub-clustering of cells identified as ANT neurons, *n* = 5535 neurons (**c**), or ACC neurons (**e**), *n* = 5671 neurons, colored by cluster number. **d, f**, Expression of marker genes for anterior/posterior thalamic nuclei in ANT neurons (**d**), or cortical layers in ACC neurons (**f**). **g, i**, Heatmap of z-scored expression of DEGs in each condition in ANT (**g**) or ACC (**i**) across time points. Columns are DEGs, rows are time points, *n* = 53-164 DEGs per time point. **h, j**, Wasserstein distances of DEGs between HR or LR to early-training in ANT (**h**), and to recent retrieval in ACC (**j**), \*\*\**P* < 0.0001 for R2, R8 and R15 HR vs LR in ANT, \*\*\**P* < 0.0001 for R8, R15 and R20 HR vs LR in ACC, One-way ANOVA with Bonferroni correction. **k, n**, Gene ontology (GO) analysis of DEGs in HR on mid-retrieval in ANT (**k**) or ACC neurons (**n**). **l, o**, Volcano plots of DEGs between HR vs LR on mid-retrieval, and remote-retrieval in ANT (**l**) and mid-retrieval and remote-retrieval day 20 in ACC (**o**). Labeled dots represent genes contributing to plasticity or histone methylation GO modules, Pvalue cutoff = 0.05, logFC cutoff = 0.015. **m, p**, Ridge plots of average expression of plasticity or histone methylation GO modules in ANT (**m**) or ACC (**p**) across retrieval days in HR (purple) or LR (green) neurons, *n* = 18-25 genes per module.

To determine whether the transcriptomic profiles of neurons associated with the HR vs LR contexts diverge over time, we used the MAST algorithm^40^ to identify differentially expressed genes (DEGs) between HR and LR cohorts across time points (Fig. 2g, 2i). We used the obtained DEGs to measure a holistic transcriptomic distance between the earliest collected time point and each successive time point using Wasserstein distance^41^. We found that the transcriptional divergence between HR and LR begins as early as recent recall, far preceding behavioral divergence, and progressively increases in both the ANT and ACC over time. While divergence in the ANT appears to attenuate by remote-retrieval, the ACC exhibits a sustained and monotonic transcriptional separation through the remote time point (Fig. 2h, 2j). This diverging pattern between HR and LR remained consistent when the Wasserstein distances were calculated using the whole transcriptome in place of just the DEGs, suggesting that a subset of highly variable genes (DEGs) may drive global divergence. Importantly, no divergence is observed when using a random subset of genes of similar size as the DEGs (extended data fig. 3a).

**Fig. 3:**
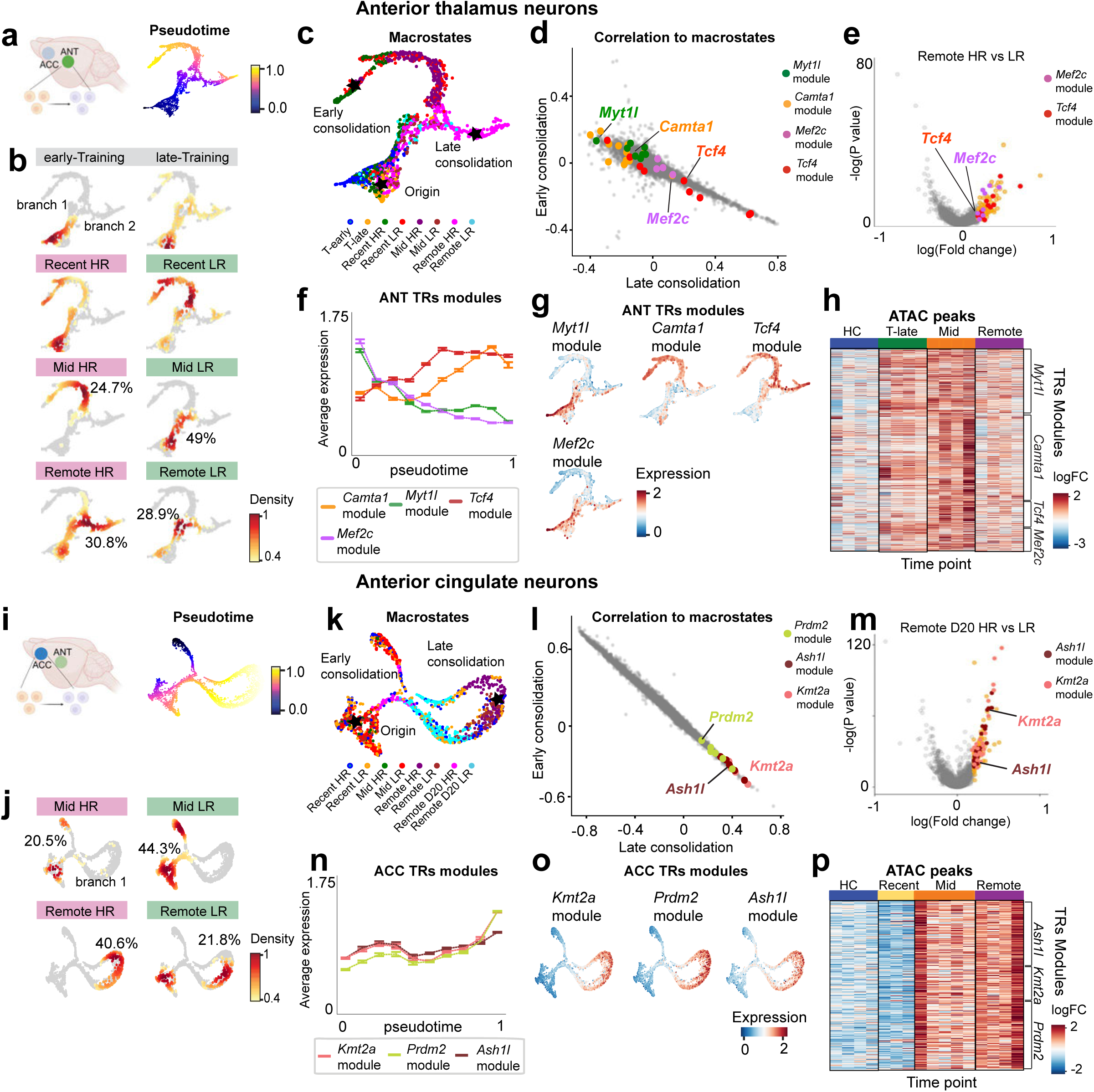
Transcriptional regulators define entry into phenotypic states associated with memory persistence. **a, i**, tSNE visualization of ANT (**a**) or ACC (**i**) neurons ordered and colored by pseudo-time. **b, j**, Density plots across training and retrieval days of HR and LR ANT (**b**) or ACC neurons (**j**) on the tSNE pseudo-time space. Percents indicate neurons with density > 0.6. **c, k**, Pseudo-time tSNE visualization of neurons from ANT (**c**) or ACC (**k**) and colored by behavioral time point. Stars highlight “apex” macrostates inferred by Palantir, which represent the origin of the pseudo-time trajectory, early or late consolidation states. **d, l**, Correlation plots of genes towards the early versus late consolidation macrostate branches for the ANT trajectory (**d**) or the ACC trajectory (**l**). Top correlated TRs with either early or late consolidation states are labeled as well as representative genes from their predicted target gene modules in matching colors. **e, m**, Volcano plots of DEGs between HR and LR at remote retrieval in ANT (e) or in ACC (m). Labeled genes are TRs’ target modules. **f, n**, Average expression of ANT or ACC TRs’ modules in ANT neurons (**f**) or ACC neurons (**n**) respectively, along the ANT or ACC tSNE pseudo-time trajectory, *n* = 13-46 genes per module. Data are mean (solid line) ± SEM (error bars). **g, o**, tSNE visualizations of expression levels of ANT (**g**) or ACC (**o**) TRs’ predicted target gene modules. **h, p**, Z-scored accessibility of ATAC peaks of ANT TRs’ modules (**h**) or ACC TRs’ modules (**p**) across time points. Columns are biological replicates, *n* = 3-5 mice per time point, and rows are ATAC peaks for TRs’ modules.

We next assessed the function of gene programs enriched in the HR condition in ANT and ACC. Gene ontology (GO) pathway analysis on the previously obtained DEGs revealed prominent modules related to synaptic plasticity in the ANT (driven by genes such as *snap25, kcna1 and cacna1e*) whereas in ACC, they were enriched for chromatin modifications, in particular histone methylation (driven by genes such as *kmt2e, kmt2a* and *setd5*); and importantly, these modules were consistent when using a pseudobulk approach to derive DEGs (Fig. 2k, 2n, extended data Fig. 3c). When the genes predicting the top ranked GO terms were overlaid onto a volcano plot of all DEGs, we confirmed that the synaptic plasticity and histone methylation gene modules were enriched in the HR condition at mid-retrieval (Fig. 2l, 2o). Notably, the plasticity related programs in ANT that peaked at mid-retrieval were transient and decreased in expression by remote retrieval, whereas the chromatin related modules in ACC exhibited sustained expression into remote time points (Fig. 2m, 2p and extended data 3b, 3d).

To assess whether specific cell types contribute disproportionately to the DEGs between HR and LR, we classified neurons in the ANT and ACC by major cell type and cortical layer, and followed their cell-type specific changes over time. We found that in the ANT the DEGs derived from Vglut2+ neurons closely mirrored those from the overall population, both in gene identity and enriched GO terms, including plasticity-related pathways at the recent time point and pruning-related processes at the remote time point (extended data Fig. 3g, 3h). This suggests that the observed transcriptional differences may be driven more specifically by a subset of Vglut2+ neurons. In the ACC, given that most neurons are Vglut1+ as expected, we additionally classified neurons by cortical layer identity. Interestingly, DEG analysis within layers revealed that neurons specifically from layers 2/3 and 6 exhibited upregulation of histone methyltransferase-related pathways at the remote time point (extended data Fig. 3i, 3j, 3k). This suggests a potential layer-specific chromatin remodeling, where changes in the layer 6 cortico-thalamic *long-range* plasticity may contribute to the layer 2/3 *within-region* stabilization of cortico-cortical connections.

Beyond cell-types, we hypothesized there may be specific “cell states” that drive the dynamic shifts in DEGs between HR and LR over time. To test this, we used density plots to visualize how neuronal clusters vary between experimental conditions (extended data fig. 3e)^42^. We obtained DEGs between HR and LR at mid retrieval, a time point marked by significant transcriptional divergence and behavioral decline, and found that in ANT, clusters 2 and 4 exhibited the highest overlap with overall DEGs, whereas in ACC this was cluster 0 (extended data Fig. 3f). These data suggest that that in each brain region some clusters may not only drive overall shifts in gene expression, but also represent the dominant cell states at the tested time points. Finally, beyond neurons, DEG assessment of non-neuronal cell types were also revealing. Whereas microglia exhibited relatively stable transcriptional profiles (extended data fig. 4a, 4b), astrocytes exhibited more dynamic, time-dependent transcriptional changes, particularly enriched in pathways associated with membrane potential regulation, kinase activity, and actin binding (extended data fig. 4c, 4d). Astrocytes in the ANT activated gene programs at earlier time points, while in the ACC, they displayed a marked upregulation of distinct HR DEGs at remote time points, aligning well with prior reports^43,44^.

**Fig. 4:**
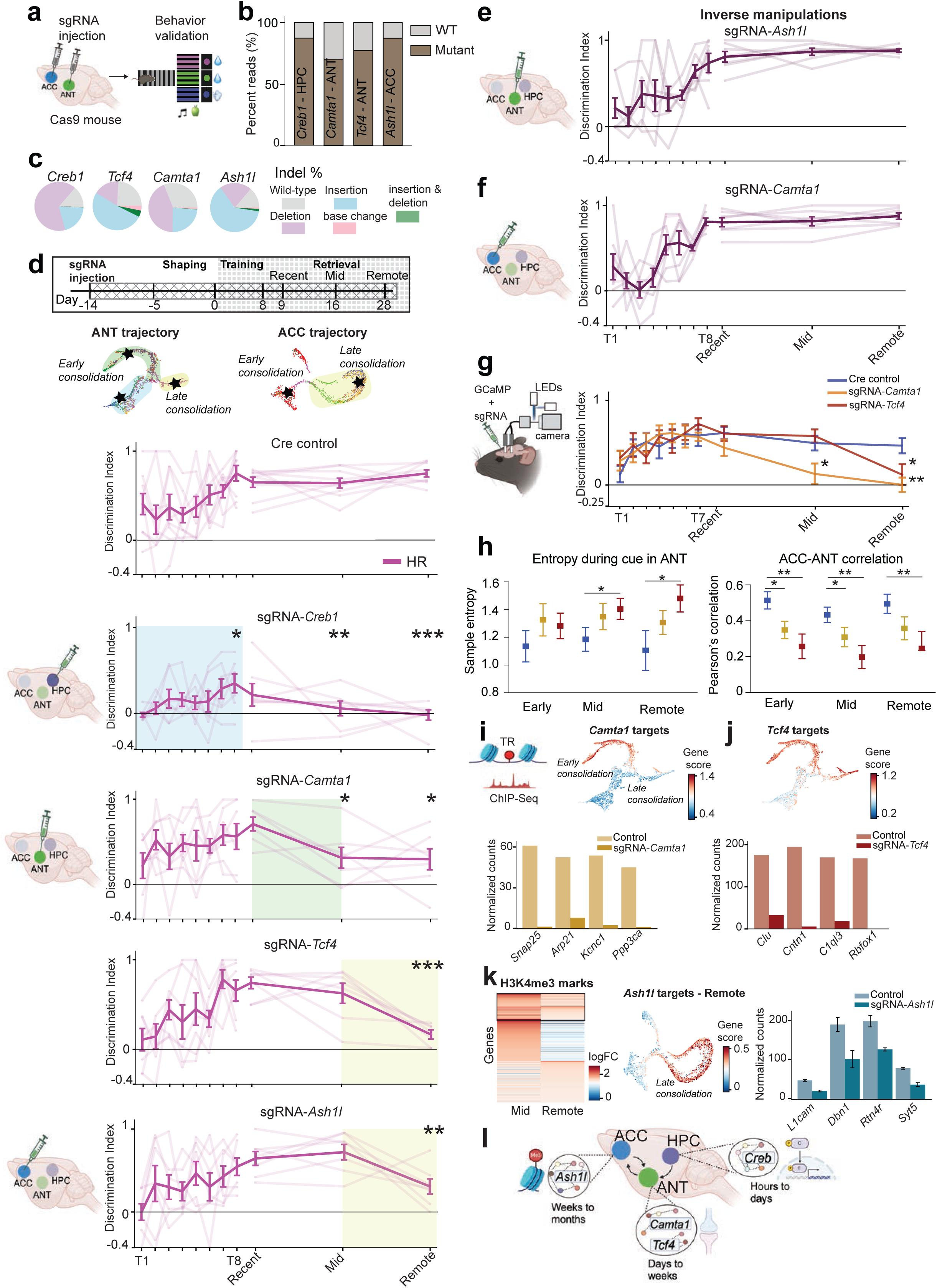
The *Camta1-Tcf4-Ash1l* thalamo-cortical transcriptional cascade is required during memory stabilization. **a**, Schematic of the CRISPR-Cas9 screen workflow. **b**, Stacked bar plot of WT vs mutant reads of amplicon regions from *Rosa26LSL-spCas9-eGFP* mice injected with AAV-sgRNA targeting the TRs shown. **c**, Breakdown of Cas9-induced indel mutations. **d**, Top: timeline of *in vivo* CRISPR-Cas9 behavioral screen, crossed pattern block represent AAV-sgRNA injection, and squared pattern block is window of behavioral testing. Below: discrimination indices for learning and recall performances in HR for mice expressing AAV-*Cre* as control or AAV-sgRNA in HPC, ANT or ACC, *n* = 7-9 mice per cohort, individual data points shown (faded lines), with mean ± SEM (solid line), \**P* = 0.048 between controls and sgRNA-*creb1* on T6, \**P* = 0.026 on T8, \*\**P* = 0.0063 on mid retrieval and \*\*\**P* = 0.0003 on remote retrieval; \**P* = 0.0288 between controls and sgRNA-*camta1* on mid retrieval and \**P* = 0.0466 on remote retrieval; \*\*\**P* = 0.0002 between controls and sgRNA-*tcf4* on remote retrieval; \*\**P* = 0.0086 between controls and sgRNA-*ash1l* on remote retrieval, One-way ANOVA with Bonferroni correction. **e, f**, Discrimination indices for mice injected with sgRNA-*camta1* in ACC (**e**) or sgRNA-*ash1l* in ANT (**f**). **g**, Discrimination indices for learning and recall performances in HR for photometry cohort, *n* = 7-10 mice, \**P =* 0.0248 between controls and sgRNA-*camta1* on mid retrieval and \*\**P* = 0.0031 on remote retrieval; \**P* = 0.0306 between controls and sgRNA-*tcf4* on remote retrieval, One-way ANOVA with Bonferroni correction. **h**, Left: sample entropy of all trials during cue of HR retrieval sessions in ANT, \**P* = 0.0361 between controls and sgRNA-*tcf4* on mid retrieval and \**P* = 0.0221 on remote retrieval. Right: pairwise Pearson’s correlations between ANT and ACC during cue of HR retrieval sessions, at ∼25 trials/mouse. mean ± SEM shown, \**P* = 0.0163 between controls and sgRNA-*camta1* on recent retrieval and \**P* = 0.0464 on mid retrieval; \*\**P* = 0.0033 between controls and sgRNA-*tcf4* on recent retrieval, \*\**P* = 0.0036 on mid retrieval and \*\**P* = 0.0011 on remote retrieval. **i, j,** Top: scored average expression of *Camta1* (**i**) *or Tcf4* (**k**) targets derived from ChIP-seq data. Bottom: bar plot of normalized counts in peak regions associated with target genes in control versus knockout samples, *n* = 1. **k,** Left: heatmap of H3K4me3 marks in ACC control samples across mid and remote retrieval. Middle: scored average expression of *Ash1l* targets derived from ChIP-seq data. Right: bar plot of normalized counts in peak regions associated with target genes in control versus *Ash1l*-knockout samples, *n* = 2. **l**, Proposed model for the role of *Camta1*, *Tcf4* and *Ash1l* in memory stabilization. Early molecular cascades, such as CREB-dependent, are triggered in the HPC and operate on the scale of hours-days, while expression of *Camta1* and *Tcf4* in the ANT extend memories beyond days through synaptic plasticity mechanisms. In the ACC, histone methylators, such as *Ash1l*, operate on longer time constants, allowing the stabilization of information across cortical ensembles.

Taken together, these results reveal that distinct transcriptional programs are activated in the ANT and ACC, which are predominantly related to synaptic plasticity and chromatin regulation respectively, and which persist to varying degrees throughout memory consolidation.^43^

## A small set of transcriptional regulators define entry into cellular macrostates associated with memory persistence

We suspected that the observed transcriptional divergence between HR and LR reflect phenotypic states that lie along a continuum, rather than in discrete clusters, from early training to late time points. Thus, to gain insight into the cell-by-cell temporal dynamics of the evolving molecular programs in the ANT and ACC we performed pseudo-time analysis on neurons from both regions (Methods). In brief, we applied the Palantir algorithm^45^, which orders cells along a continuum based on phenotypic similarity. A starting cell is first defined to orient the trajectory, and the algorithm automatically identifies multiple end states (i.e. states with highest variance), and assigns every cell with a probability of traversing into each end state. Palantir has been successfully used in multiple developmental and cancer studies for trajectory detection^46^. While such pseudo-time trajectory inference algorithms have not been traditionally used in conjunction with behavior, here we used it to capture the temporal progression of cellular states associated with memory persistence, and validated this approach using multiple algorithms, and mapping of expected biological changes, as detailed below.

In our implementation of the Palantir algorithm, we seeded a random cell from the earliest time point (early-training in ANT) as the start of the pseudo-time trajectory and ordered all training, HR and LR remaining cells along a pseudo-time continuum based on phenotypic similarity (Fig. 3a). In the ANT, Palantir identified two apex branches (or most extreme phenotypic states): a leftmost “branch 1” and a rightmost “branch 2”. We next overlaid all cells from the HR and LR conditions onto this trajectory, and found atypical distribution of cells from each timepoint. Neurons from training days occupy earlier states along the pseudo-time and by recent-retrieval, both HR and LR cells begin to progress along the leftmost “branch 1”. However, by mid-retrieval, while HR neurons maintained their phenotypic state along “branch 1”, LR neurons occupied a phenotypic state closer to the start of the trajectory. By remote retrieval, most HR neurons reached a new phenotypic state along “branch 2”, while again LR neurons remained near the origin of the pseudo-time (Fig. 3b). These data thus identify unique trajectories for HR and LR neurons during memory stabilization, where HR neurons explore three consecutive phenotypic macrostates, one which is unique to HR (representing late consolidation), and which may define memory persistence (Fig. 3c).

Notably, the observed early and late consolidation cellular macrostates were not simply batch effects related to sample collection at different time points or driven by a single mouse’s performance. This was evidenced by cell densities and branch occupancy per biological replicate which were similar to those observed from all sampled cells (extended data Fig. 5a). We also confirmed that trajectory progression is not driven by any uneven technical recovery of specific cell classes, as indicated by largely even entropy scores of major cell types and cortical layers across pseudo-time intervals (Methods; extended data fig. 5c); rather we observe segregation of cell clusters suggesting that pseudotime evolution is driven by cell states (extended data Fig. 5e left). Moreover, shifts between the HR and LR conditions along the trajectory do not merely represent passage of time, but rather changes in cell states. For instance, not all ANT HR cells reached the early or late consolidation macrostates (branch 1 or branch 2), rather ∼30% of HR cells reached these states (comparable to the ∼20% of cells that encode the HR condition during in vivo imaging^45^), further underscoring that only a subset of neurons are behaviorally relevant and undergo consolidation related transcriptomic changes (Fig. 3b). Relatedly, by mapping *Fos*+ neurons onto this pseudo-time (∼30% of all neurons per time point in our dataset, extended data fig. 5f left), we again observed similar HR vs LR trajectory shifts that mirrored the shifts that we observed for the entire sampled population (extended data fig. 5g). Of note, we found a large overlap between DEGs between HR and LR from *Fos*+ cells and all DEGs, indicating that *Fos*+ cells represent a highly relevant population (though are not the sole contributors) to the observed transcriptional differences between HR and LR (extended data fig. 5i, left).

To further validate shifts in phenotypic states, we performed unbiased grouping of cells into “neighborhoods” and asked whether those neighborhoods were enriched in HR or LR neurons. To do so, we used MiloR^47^, which allows for the statistical assessment of cellular abundances by partitioning the space into overlapping neighborhoods of cells, and importantly, by constructing such neighborhoods in the original high-dimensional phenotypic space. Such abundance analysis further confirmed that HR cells were most enriched in the trajectory apexes representing the “late” consolidation state (extended data fig. 6a left). Additionally, when examining the DEGs from the earliest time point, early-training, we observed that the highest expression was in neurons closer to the origin of the pseudo-time, while HR DEGs from remote-retrieval were most highly expressed on the “late consolidation branch” (extended data fig. 6b, left). To validate our trajectory methodologies on existing public datasets, we mapped molecular programs that have previously been reported to occur during learning and early consolidation (learning-associated IEGs such as *Fos*, *Junb*, *Homer1*, *Egr1*, *Erg2*, *Egr3*, *Egr4*)) and found that they were most highly expressed by neurons occupying phenotypic states closer to the start of the pseudo-time and the early consolidation macrostate (extended data fig. 6c). In addition to the Palantir algorithm, we also applied the CellRank^48^ algorithm as an independent verification. This algorithm infers the start and end points of the pseudo-time in an unbiased manner (Methods), and again, we observed the identification of two distinct cellular macrostates (representing early and late consolidation) associated with memory persistence (extended data fig. 6d left, and fig. 6e-g). Finally, in a separate cohort of mice, we also validated these evolving gene expression programs using qPCR showing an expected upregulation of plasticity related genes at early retrieval and structural related genes at remote retrieval in HR mice (extended data Fig. 7d). Taken together, these results demonstrate that we are capturing phenotypic macrostates associated with memory persistence that are technically robust and biologically meaningful.

We repeated the above pseudo-time analyses for the ACC neurons (Fig.3i). Given ACC is known to have a late role in memory consolidation^43,46^, we included samples from recent retrieval as the earliest time point. We seeded a random cell from recent retrieval as the start of the ACC pseudo-time. We interpret this as meaning that taking our earliest sampled time point as origin may not accurately represent a true baseline state, but rather somewhere on the continuum of ongoing consolidation. In ACC neurons, Palantir identified one apex macrostate (branch 1). We again overlaid the cell densities from HR and LR conditions per time point (and across animal replicates) and found that HR cells were denser on “branch 1” (∼40%) unlike LR neurons which occupied both “branch 1” (∼22%) and the area near the origin (Fig. 3j, extended data fig 5b, 6a right). We noticed that ACC neurons alternated between the “origin” and “branch 1” more than once (extended data fig. 5j), suggesting that ACC neurons may undergo dynamic and reversible transitions between these two states, leading to sustained transcriptional activation. We confirmed that trajectory progression was not driven by anatomically defined cell classes (Methods; extended data Fig. 5d, 5e right). *Fos*⁺ cells exhibited transcriptional shifts similar to those of the overall neuronal population (extended data Fig. 5f right, 5h). Additionally, early and late differentially expressed genes mapped specifically to the early and late branches of the trajectory, respectively (extended data Fig. 6b, right). As an additional validation of our pseudo-time approach, we performed trajectory inference on V1 visual cortex neurons from mice trained in the VR task. The Palantir algorithm did not identify apex branches, nor did it reveal divergence between HR and LR conditions at remote retrieval, suggesting that neurons not involved in memory processing do not undergo significant cell state transitions (extended data Fig. 6j-6l).

Since transcriptional regulators (TRs) have been previously shown to be key drivers of branch commitment^45,48^, we analyzed TRs whose expression changes correlated with early and late branches of the ANT and ACC trajectories. We computed the spearman’s correlation between gene expression levels and “macrostate” probabilities (Methods), and derived a list of candidate TRs that drive branch bifurcations (Table S3). In the ANT, we considered the top four such TRs whose expression correlated with early consolidation (*Camta1*, M*yt1l*) and late consolidation (M*ef2c* and T*cf4*) macrostates respectively (Fig. 3d). Strikingly, the predicted downstream target genes of these TRs comprised a large fraction (20-30%) of the total DEGs representing each branch (Fig. 3e), and whose average expression of TRs’ module covaried with branch states (Methods, Fig. 3f, 3g and extended data fig. 6h, 6i), thus strongly supporting a role in branch commitment. We verified that these TRs are well expressed in ANT (extended data fig. 7a, 7c) and subsequent ATAC-seq analysis further confirmed increased accessibility of these four TRs by day mid-retrieval, with varying levels of persistence that mirrored the expression pattern of these TRs (Fig. 3h; extended data fig. 8c-8e, 8g left). These data thus identify a small set of key TR-candidate genes that appear to orchestrate global molecular programs enabling entry into early and late consolidation macrostates in ANT.

In the ACC, using a similar set of analyses, we identified three TRs that defined the late consolidation macrostate (*Ash1l*, *Kmt2a* and *Prdm2*), all of which have a role in histone methylation (Fig. 3l-3m and extended data fig. 6h, 6i). While chromatin modifications, such as histone methylation, histone acetylation, and DNA methylation have all been described as potential mechanisms that support memory consolidation, the identified TRs strikingly favored histone methylation over other modifications as critical for long term cortical consolidation. These TRs are well expressed in the ACC (extended data Fig. 7b, 7c), and that while their ATAC peaks were most accessible on mid-retrieval (returning to baseline by remote-retrieval, extended data fig. 8g right), the peaks for their associated modules remained accessible through remote time points (Fig. 3p). These data reveal sustained expression of chromatin remodeling programs enriched in ACC in the late phase of consolidation.

Interestingly, we observed that a large fraction of DEGs emerging in late-training in ANT appear to re-activate during later recall (Fig. 2g), suggesting that some of the molecular programs defining memory persistence (associated with HR recall at remote time) are activated quite early, even at the time of learning, well before behavioral divergence. The early engagement of these transcriptional programs, prior to observable behavioral divergence between HR and LR conditions, raises the possibility that successful memory consolidation relies on the timely recruitment of specific gene networks at the moment of learning. These early transcriptional signatures may serve as predictive markers of future memory strength or durability.

Taken together, these combined pseudo-time results indicate that the ANT and ACC neurons may recruit a sparse and distinct set of transcriptional regulators, which turn on broad transcriptional programs over different time scales, that ultimately shape macrostates associated with memory persistence. The ANT recruits an early activation of plasticity-related programs, potentially driven by *Camta1, Myt1l, Mef2c and Tcf4*, whereas the ACC recruits predominantly histone methylators, such as *Ash1l*, *Kmt2* and *Prdm2,* whose activity remain elevated longer, potentially conferring more enduring effects on memory storage in cortical circuits.

## A Camta-Tcf4-Ash1L thalamo-cortical transcriptional cascade is required for memory stabilization

To test whether the identified TRs have causal roles in memory stabilization, we next performed region-specific knockout of these genes using CRISPR-based *in vivo* manipulations. We tested *Camta1*, *Myt1l*, *Mef2c* and *Tcf4* in ANT; *Kmt2a* and *Ash1l* in ACC; we also included *Creb1* in HPC as a positive control. We began by screening the efficiency of many single guide RNAs (sgRNAs) *in vitro* (∼8-10/gene) by transfecting them into Neuro2A cells and using Sanger-based analysis to analyze the editing efficiency of collected DNA (extended data fig. 9a, 9b, Methods). We then selected the most efficient single sgRNA per gene (with high predicted on-target efficiency and minimal predicted off-target effects) and producing mutant reads greater than 50% (extended data fig. 9c, 9d), and packaged these into an adeno-associated virus (AAV) together with Cre-eGFP, for in vivo behavioral testing. We delivered AAV9-U6-sgRNA-hSyn-Cre or a control AAV9-Cre into either the ANT or the ACC of Rosa26LSL-spCas9-eGFP mice^49^, which express Cas9 in a Cre-dependent manner (Fig 4a). We verified the successful knockout of targeted genes *in vivo*, and catalogued the type of mutations that occurred, by performing Next Generation sequencing (NGS) (Fig. 4b). We confirmed knockout at the protein level by performing immunostaining on brain slices (extended data Fig. 9e-9g).

We allowed animals to recover and to express the construct for two weeks before starting behavioral testing (Fig. 4d top). Control Rosa26LSL-spCas9-eGFP mice expressing only Cas9 (and Cre, but without sgRNA) successfully learned the task, as demonstrated by high DI’s across time (Fig. 4d,). Mice with knockout of *Creb1* in hippocampus displayed, as expected, strong learning and early memory deficits (Fig. 4d). Interestingly, mice expressing sgRNAs targeting the ANT or ACC TRs displayed no observable learning deficits. However, knockout of a subset of those TRs led to striking and temporally restricted consolidation deficits that were specific to ANT (early to mid-consolidation) or ACC (late consolidation). Knockout of C*amta1* resulted in impaired recall at the intermediate time point, whereas knockout of T*cf4* produced a recall deficit only at remote time (Fig. 4d, *P* =0.0288 between controls and sgRNA-*Camta1* on mid retrieval, *P*=0.0002 between controls and sgRNA-*Tcf4* on remote retrieval, One-way ANOVA with Bonferroni correction). Knockout of *Myt1l* or *Mef2c*, on the other hand, resulted in no significant memory impairments (extended data fig. 9h).

Of the genes tested in ACC, only knockout of *Ash1l*, resulted in an isolated remote memory deficit, leaving both learning and recent memory intact (fig. 4d, *P*=0.0086 between controls and sgRNA-*ash1l* on remote retrieval, Bonferroni’s multiple comparison correction). Knockout of *Kmt2a* resulted in a trending but not significant remote impairment, which might be explained by redundancy of regulators from the same family (i.e. Kmt2c, extended data fig. 9h). Strikingly, the functional effects of *Camta1* in ANT and *Ash1l* in ACC were region-specific, as inverse manipulations, i.e., knockout of *Camta1* in ACC or *Ash1l* in ANT produced no behavioral deficits (Fig.4e, 4f).

To explore how these TRs and their associated molecular programs may give rise to functional changes in the ANT-ACC circuit, we performed GCaMP-based neural-activity recordings in ANT and ACC longitudinally during behavior, comparing controls with *Camta1* or *Tcf4* KO (extended data Fig. 10a; Methods). Notably, all three cohorts of mice learned equally well, but again we were able to robustly replicate that the *Camta1*-KO have early consolidation deficits whereas the *Tcf4*-KO have late consolidation deficits (Fig. 4g). Furthermore, in the neural data, we found that the *Camta1* and *Tcf4* KO display early increases in neural entropy (a measure of neural variability or signal irregularity) together with significant deficits in ANT-to-ACC functional correlations (reflecting weaker long-range communication throughout the consolidation window) that precede behavioral deficits in memory maintenance (Fig. 4h). While higher resolution imaging in future studies will be more informative, these initial data suggest that *Camta1* and *Tcf4* may coordinate physiologic changes in ANT that coordinate ANT-ACC functional connectivity and plasticity necessary for memory stabilization.

Finally, to reveal mechanistic insights into the transcriptional targets of *Camta1*, *Tcf4*, and *Ash1l* and their roles in memory consolidation, we performed ChIP-seq analyses during behavior (extended data Fig. 10b). We found marked depletion of signal in *Camta1*- and *Tcf4* KO cohorts (extended data Fig. 10c). By comparing controls and KOs, we confirmed that the most strongly depleted peaks in *Camta1* knockout mice were associated with genes involved in voltage-gated channel activity and plasticity related pathways (extended data Fig. 10d top, 10e top), whereas *Tcf4* knockout depleted peaks were in genes associated with cell adhesion (extended data Fig. 10d bottom, 10e bottom), Strikingly, when we overlaid the identified *Camta1*- and *Tcf4*-target genes onto the ANT trajectory, we found that their expression was elevated primarily along the early (Fig. 4i) or late consolidation branch respectively (Fig. 4j). Meanwhile, for ACC, we examined the changes in H3K4me3 methylation marks across time, as well as how Ash1l KO might disrupt this. We found an interesting temporal shift in methylation of gene programs over time—from plasticity related pathways in mid-retrieval to structural components at remote retrieval (Fig. 4k left, extended data Fig. 10f). Reassuringly, *Ash1l* knockout resulted in depletion of such plasticity-associated genes at mid-retrieval, whereas by remote retrieval, as expected, the most depleted peaks corresponded to genes involved in structural regulation (Fig. 4k, extended data Fig. 10g, 10h).

In sum, by identifying the transcription factors *Camta1* and *Tcf4* in the ANT, and the histone modifier *Ash1l* in the ACC, that have sequential, circuit-specific, and causal contributions to memory maintenance, we provide a mechanism for the continuous stabilization of memory from days to weeks (Fig. 4l).

## Discussion

The earliest discovered signaling molecules important for learning and memory were components of the calmodulin/cAMP pathway^50–52^. In addition to these transient mechanisms, the role of protein synthesis, and in particular the activation of the cAMP dependent transcription factor CREB1, was found to be important for enabling longer lasting forms of synaptic plasticity on the order of hours to days. Still, the molecular programs underlying the maintenance of memories on longer time-scales have remained elusive. Here, we expand our molecular understanding of memory beyond the well-studied hippocampus and identify distinct transcriptional regulators, operating initially in the anterior thalamus (*Camta1* and *Tcf4*), then in the anterior cingulate cortex (*Ash1l*), that propagate memory maintenance progressively from days to weeks. Notably, these TRs were not required during learning, but instead had defined, sequential, time-limited roles in memory maintenance. Additionally, these effects were notably circuit specific, where *Camta1* and *Tcf4* had critical functions in the anterior thalamus, and *Ash1l* in ACC, but not vice versa. Furthermore, we observed little to no effects on memory maintenance when manipulating other selected TRs, such as *Myt1l*, *Mef2c* or *Kmt2a,* in the thalamo-cortical circuit, whose gene-modules also co-varied on similar time-scales throughout behavioral task, further underscoring a critical causal role for *Camta1, Tcf4* and *Ash1l* in orchestrating progressively longer time-scale memory stabilization.

These results thus highlight several important aspects of the memory stabilization process: 1. That beyond the hippocampus, the thalamo-cortical circuit has a critical contribution to memory stabilization, 2. That memory stabilization is an active and continuous process requiring the successive recruitment of transcriptional programs operating on progressively longer time-scales, and 3. These time-limited transcriptional programs operate in a circuit-specific manner, providing an explanation for why multiple circuits across the brain are recruited to support continuous memory stabilization.

We observed that the ANT and ACC engage distinct functional pathways to support memory stabilization. In the ANT, we found significant enrichment of plasticity-related programs. For instance, the calmodulin/cAMP-sensitive transcription factor Camta1 and the basic helix-loop-helix transcription factor Tcf4 both have activity dependent gene-modulatory roles, and regulate neuronal excitability and synaptic plasticity^53–56^. Conversely, the ACC preferentially recruits chromatin remodeling programs and histone methylation emerges as a critical mechanism for extending memory persistence. Many downstream targets of ASH1L^59^ are components of the synapse known to be involved in synapse stability, including transmembrane proteins, adhesion molecules, and structural filaments^60,61^. *Ash1l* has also been implicated in activity-dependent synaptic pruning^62^, a process that refines neural circuits and contributes to the long-term maintenance of memories. In human studies, *CAMTA1*, *TCF4*, and *ASH1L* have all been linked to intellectual disability or memory impairment^63–65^.

It is interesting to note that while Ash1l emerges as a late phase regulator of memory stabilization in the ACC, its targets are largely overlapping with those of Camta1 and Tcf4 in the ANT, targeting synaptic plasticity and adhesion/structure respectively. This suggests that Ash1l methylation in ACC may function as a mechanism to “prime” ^57,58^ and prolong plasticity/structural gene targets necessary for extending synaptic and circuit time-scales. Indeed, histone methylation marks tend to persist longer than transcription factor activation, aligning with a model where the sequential recruitment of transcriptional regulators across distinct brain regions enable a progressive reorganization of memories from transiently plastic states to more lasting representations.

The broad regulatory scope of transcriptional regulators presents a significant challenge in distinguishing memory-specific effects from broader alterations in cellular state. To better contextualize the functional relevance of CAMTA1, TCF4, and ASH1L in memory consolidation, we incorporated multiple complementary approaches that strengthen the link between these TRs and their role in supporting memory. First, the *in vivo* functional recordings demonstrated that *Camta1* and *Tcf4* knockout result in increased neural variability together with significant deficits in ANT-to-ACC functional correlations, thus reflecting weaker long-range communication throughout the consolidation window. Then, ChIP-seq profiling of CAMTA1 revealed that it predominantly binds genes associated with synaptic plasticity, while TCF4 targets are enriched for genes involved in structural regulation. Profiling of H3K4me3 marks in ASH1L-KO suggests that ASH1L regulates both plasticity and structural-related programs. These findings suggest that each factor contributes to distinct transcriptional modules that support different phases or components of memory consolidation. While these results indicate that CAMTA1, TCF4, and ASH1L help coordinate specific transcriptional programs underlying memory-associated state transitions, future studies using more temporally and spatially resolved or graded manipulations—such as CRISPRa/i will be critical to more precisely link changing molecular programs to physiology and behavior.

These findings, and work from other labs, collectively support the notion that the basic biological substrates that convert transient stimuli into long-lasting cellular states that maintain cell identity^66,67^, or behavioral states and longevity^68–71^ can be co-opted to support long-lasting memories. For instance, emergent network properties can push transcriptional programs into long-lasting states, which can operate on different time-scales^72^. Relatedly, epigenetic programs are known to support even more lasting cellular phenotypes, and pharmacological studies have already highlighted important roles in adult brain during learning and memory^73,74^. Here by identifying specific TRs that have sequential contributions to memory maintenance, we provide a mechanism for the continuous stabilization of memory over prolonged time-scales.

## Acknowledgements

We thank the Rockefeller Flow Cytometry Resource Center for sorting of samples, and the Single Cell Analytics Innovation Lab (SAIL) at Memorial Sloan Kettering Cancer Center for sequencing and processing scRNA-seq samples. We thank H. Tan in the laboratory of J. Friedman for generously providing Rosa26LSL-spCas9-eGFP mice; Z. Gershon for help troubleshooting single cell dissociations; E. Azizi, O. V. Goldman and A. Sziraki for helpful discussions related to analysis of scRNA sequencing data; N. Blobel and S. Nakandakari for assistance with *in vitro* CRISPR-based manipulations and sharing reagents; Y. Kishi for sharing the ATAC sequencing protocol; N. Heintz, E. Azizi and members of the Rajasethupathy Lab for comments on the manuscript. This work is supported by a Kavli Neuroscience Institute pilot grant from the Rockefeller University and a graduate fellowship from the Kavli Neural Systems Institute (A.T), and grants from Irma T. Hirschl / Weill Caulier Trust and the National Institutes of Health under award numbers DP2AG058487 and RF1NS132047 (P.R).

## Author Contributions

A.T, C.C and P.R. conceived the study and designed the experiments. A.T performed the optogenetic experiments, collected scRNA sequencing samples with assistance from M.G, and developed the analysis pipeline for scRNA sequencing. R.S assisted with scRNA-sequencing analysis and pseudo-time analysis. A.T and C.C developed a pipeline for *in vitro* and *in vivo* validation of sgRNAs, performed *in vivo* Crispr-manipulations and behavioral testing. T.E performed photometry experiment and analysis. C.C performed qPCR and immunohistochemistry/Western Blot validations. Y.H collected samples for ATAC-sequencing and R.K analyzed the ATAC-sequencing data. C.C collected samples for ChIP-sequencing, P.J.H processed samples, R.K analyzed the ChIP-sequencing data, A.T performed downstream analysis. A.T, C.C and P.R. wrote the manuscript with input from all authors. P.R. supervised all aspects of the work.

## Methods

### Mice

All mice were purchased from The Jackson Laboratory. Six-to eight-week-old wild-type C57Bl6/J male or female mice (Jackson Laboratories, cat. no. 000664) were group-housed three to five in a cage with unlimited access to food and water, unless mice were water restricted for behavioral assays, in which mice were given a total of 1 mL of water per day. For CRISPR experiments, six- to eight-week-old male or female Rosa26-LSL-Cas9 knockin mice (Jackson Laboratories, cat. no. 024857) were housed under the same conditions. All procedures were done in accordance with guidelines approved by the Institutional Animal Care and Use Committees (protocol no. #22087H) at The Rockefeller University. Number of mice used for each experiment was determined based on expected variance and effect size from previous studies and no statistical method was used to predetermine sample size.

### Surgeries

All surgical procedures and viral injections were carried out under protocols approved by the Rockefeller University IACUC. Mice were anesthetized with 1-2% isofluorane for the entire duration of the procedure and positioned on a Kopf stereotactic apparatus with a heating pad. Puralube vet ointment was applied to the eyes to prevent drying and 0.2 mg/kg meloxicam was administered intraperitoneally using a 1mL syringe and 23G needle. Hair from the scalp was trimmed, and the area was sterilized povidone-iodine and ethanol. A midline incision was made with a sterile scalpel and holes for injection sites were made using a sterile 0.5 mm micro drill burr (Fine Science Tools) through the skull. All virus was injected using a 24G beveled needle (World Precision Instruments) in a 10ul NanoFil Sub-Microliter Injection syringe (World Precision Instruments) controlled by an injection pump (Harvard Apparatus) at a rate of 100 nL/min. Following viral injection, the needle was raised 0.1 mm above the injection site for 3 minutes (to prevent backflow) before slowly raising the needle out of the skull. 4-0 vicryl and Vetbond (3M) was used to close the incision. For mice used for head-fixed behavior, a custom titanium headplate was adhered to the skull with Metabond. For mice that required cannulas, the cannulas were implanted immediately following viral injection. Animals were allowed to recover on a heating pad for 1 hour and given meloxicam tablets. Following viral injections, mice were kept for 3 weeks to allow for adequate expression of the viral construct before behavioral testing or histology.

### Viral injections

- Coordinates used were: CA1 (A/P: − 1.5 mm, M/L: ±1.5 mm, D/V: −1.5 mm); ACC (A/P: +1.0 mm, M/L: ±0.35 mm, D/V: −1.4 mm); ANT (A/P: − 0.85 mm, M/L: ±0.6 mm, D/V: −3.55 mm); EC (A/P: −4 mm, M/L: ±3.75 mm, D/V: −4.2 mm); RSC (A/P: −2 mm, M/L: ±0.4 mm, D/V: −0.5/−0.8 mm); BLA (A/P: −1.3 mm, M/L: ±3 mm, D/V: −4.5 mm).
- In stGtACR2 inhibition experiments, 900nl of rgAAV-hSyn-Cre (Addgene, cat. no. 105553, 1.3 × 10^13^ vg/mL) was injected bilaterally in ACC and 900nl of AAV1-hSyn1-SIO-stGtACR2 (Addgene, cat. no. 105677) was injected bilaterally in ANT, EC, BLA, RSC. pAAV-CKIIa-stGtACR2-FusionRed (Addgene, cat. no. 105669) was injected bilaterally in ACC or HPC (AP: -1.5mm, ML: ± 1.5mm, DV: - 1.5mm).
- In CRISPR-Cas9 knockout experiments, 600nl of AAV9:ITR-U6-sgRNA-hSyn-Cre-2A-EGFP (Addgene, cat. no. 60231, 1.0 × 10^12^ vg/mL) was injected bilaterally in ANT or 900nl was injected bilaterally in ACC. For experimental controls, 600nl of AAV9-CRE (1.0 × 10^11^ vg/mL) was injected bilaterally in ANT or ACC.
- In the photometry experiments, Rosa26-LSL-Cas9 control mice were injected with 800nl AAV9-CAMKIIa-jGCaMP8m (Addgene, cat. no. 176751-AAV9, 5 × 10^12^vg/mL) contralaterally in ANT (A/P: −0.85, M/L: ±0.6, D/V: −3.55), and ACC. Rosa26-LSL-Cas9 mice for knockout testing were additionally injected with 500nl AAV9:ITR-U6-sgRNA-hSyn-Cre-2A-EGFP (Addgene, cat. no. 60231, 1.0 × 10^12^ vg/mL) bilaterally in ANT.

### Cannula implants

For optogenetics, surgeries were carried out as previously described. Immediately after viral injection, animals were implanted with fiber optic cannulas (Doric Lenses). Mice were implanted bilaterally with 200µm diameter cannulas (0.22 NA, Doric Lenses). Cannula implants were slowly lowered using a stereotaxic cannula holder (Doric) at a rate of .001 mm/sec reaching 0.2 mm dorsal to the injection site. Throughout the implantation procedure, the injection area was continually flushed with 0.9% saline and suctioned. Optic glue (Edmund Optics) was then used to secure the cannula to the skull surface, and a custom titanium headplate was affixed as previously described.

For photometry, mice were contralaterally implanted with 1.25 mm ferrule-coupled optical fibers (0.48 NA, 400 mm diameter, Doric Lenses) cut to the desired length so that the implantation site is ∼0.2 mm dorsal to the injection site in ANT and ACC.

### Optogenetic manipulations

#### Optogenetic inhibition

For projection specific expression of stGtACR2, mice were injected with rgAAV-hSyn-Cre (Addgene, cat. no. 105553) bilaterally in ACC and AAV1-hSyn1-SIO-stGtACR2 (Addgene, cat. no. 105677) bilaterally in the ANT, BLA, EC or RSC. For local inhibition, AA9-CKIIa-stGtACR2-FusionRed (Addgene, cat. no. 105669) was injected bilaterally in ACC or HPC. Control cohorts were injected with rgAAV-hSyn-Cre (Addgene, cat. no. 172221) in ACC and AAV9-hSyn-mCherry (Addgene, cat. no. 114472). Volumes and titers are described previously. A blue 470 nm light was delivered during the cue zones of training sessions (at a power of 15 mW measured at the fiber tip). No inhibition was carried out during retrieval sessions.

### Histology

Mice were transcardially perfused with 20 mL cold PBS and 20 mL cold 4% paraformaldehyde (in PBS). Brains were submerged in 4% paraformaldehyde at 4 °C overnight. The next day, brains were submerged in a 30% sucrose (dissolved in PBS) for 24 hours at 4 °C. For histology, brains were sliced into 60-μm coronal sections using a freezing microtome (Leica SM2010R) and stored in 1x PBS. For immunostaining, fixed brain sections were blocked in solution of 3% normal donkey serum, 5% BSA, and 0.2% Triton X-100 in 1× PBS for ∼3 h and incubated with primary antibody overnight at 4 °C. Sections were washed 3 times in PBS and incubated in the appropriate secondary antibody for ∼2.5 h at RT. Following 3x 5 min washes in PBS, free-floating sections were stained with DAPI (1:1,000 in PBST), and mounted on slides with ProLong Diamond Antifade Mountant (Invitrogen). Images were acquired at 10x and 20x magnification with a Nikon Inverted Microscope (Nikon Eclipse Ti). Primary antibodies include Cre Recombinase (D7L7L) XP Rabbit mAb (Cell Signaling Technology, cat. no. 15036, 1:100 dilution), Creb (48H2) Rabbit mAb (Cell Signaling Technology, cat. no. 9197, 1:50 dilution), anti-Camta1 Rabbit pAb (Millipore Sigma, cat. no. SAB4301068, 1:100 dilution). Secondary antibodies include Alexa Fluor 647-conjugated AffiniPure donkey anti-rabbit IgG (Jackson ImmunoResearch, cat. no. 711-605-152, 1:250 dilution) and AlexaFluor 647-conjugated AffiniPure donkey anti-mouse (Jackson ImmunoResearch, cat. no. 715-606-151).

### Behavior

#### Virtual reality system

For the behavioral experiments, we used a custom-built virtual reality environment by adapting a previously reported task^73^. Briefly, a 200-mm-diameter styrofoam ball was axially fixed with a 6-mm-diameter assembly rod (Thorlabs) passing through the center of the ball and resting on 90° post holders (Thorlabs) at each end, allowing free forward and backward rotation of the ball. Mice were head-fixed in place above the center of the ball using a headplate mount (Thorlabas). Virtual environments were designed in the virtual reality MATLAB engine ViRMEn. The virtual environment was back-projected (Kodak Ultra Mini Portable Projector) onto white fabric stretched over a clear acrylic hemisphere with a 14-inch diameter placed ∼20 cm in front of the center of the mouse, encompassing ∼220° of the mouse’s field of view. The rotation of the styrofoam ball was recorded by an optical computer mouse (Logitech) that interfaced with ViRMEn to transport the mouse through the virtual reality environment. A National Instruments Data Acquisition (NIDAQ) device was used to record lick events (as capacitance changes on the lick port), and to trigger the various Arduinos controlling tones, odors, air puff, as well as optogenetic stimuli.

#### Behavioral shaping

Mice were put on a restricted water schedule, receiving 1 mL of water per mouse on a given day. Body weight was monitored daily to ensure it was maintained above 80% of the pre-restriction measurement. After 3 days of water deprivation, mice were habituated to the Styrofoam ball for 5 days by receiving their 1 mL of water per day while head-fixed, and exposed to a linear track and a cue zone, where an auditory cue was delivered (Hz), and an outcome zone where they received 5 s of water delivery. If a mouse did not drink 1 mL of water, it was supplemented with water that day to a total of 1 mL. After 5 days of the shaping protocol, training began.

#### Behavioral training

Each trial was initiated in a neutral track zone. Next, mice were transported to a cue zone where they learned to use contextual presented cues to predict the paired reward (water) or punishment (air puff to snout) in the outcome zone. The contextual cues consisted of visual cues (colors and shapes on the walls of the track), auditory cues (outputted by a thin plastic speaker (Adafruit), and olfactory cues (released from a custom-built olfactometer). The visual cues were generated within the ViRMEN GUI, and both auditory and olfactory cues were outputted by Arduino code under the control of ViRMEN code. The contexts used were: 1. Reward-HR (yellow rectangles for visual, isoamyl acetate for odor, 5KHz tone for auditory), 2. Reward-LR (pink hexagons, benzaldehyde, 7KHz tone), 3. Aversive (blue triangles, octanal, 9.2KHz). All three contexts were interleaved and randomized on training days. The HR context appeared with ∼50% frequency, the LR context appeared with ∼22% frequency, and the aversive context appeared with ∼28% frequency.

After the cue zone, mice were transported to an outcome zone, where after reward cues, they received water if they touched the lick port. After the aversive cues, two air puffs (35 psi) were released by a solenoid (Precigenome, Isolation Valve, 20NC, 0.032” (0.8mm) Orifice, Diaphragm, 2-Way) controlling airflow into a pipette tip cut to a fine 1 cm away from the snout. Although mice could self-initiate movement on the ball, which would generate visual movement down the VR track, they were transported through the rooms on a timed schedule, regardless of the distance they ran on the ball. For a single trial, mice were transported through a neutral start track (8 seconds), cue zone (5 seconds) and outcome zone (5 seconds) in training, resulting in 40-50 trials per session. In retrieval durations slightly shorter, in neutral start zone (5 seconds), cue zone (5 seconds) and outcome zone (3 seconds) in retrieval, resulting in 20-30 trials per session.

Performance on the task was assessed by average anticipatory lick rate measured from the last 2 seconds of the cue zone immediately preceding the outcome zone. Mice were trained for at least 7 days and up until 9 days, depending on when they met criteria (DI >= 0.3). During training, reinforcement (air puff or water) was always paired to the outcome zone. In contrast, during retrieval, mice were not presented with any reinforcement in the outcome zone. Thus, during retrieval, both anticipatory licking and licking during the outcome zone were considered. For some longitudinal experiments required testing over multiple retrieval days, the mice had to be retrained on the task to avoid memory extinction and loss of engagement in the task in future timepoints. Immediately following testing on the same day, these mice received 10 minutes of a “retraining” session where reinforcement (water or puff) was re-introduced.

The overall time course of the behavior consisted of the following phases: habituation (5 days), training (T1-T9), retrieval (R1-R30). During the habituation phase, each mouse was habituated for 15 minutes. During the training phase, each mouse was trained for 15-17 minutes per session. During the retrieval phase, mice were tested for 5-6 minutes. Recent retrieval was defined by retrieval days R1 to R8 following the final day of training. Mid retrieval was defined by retrieval days R8-R15 and remote retrieval was defined by retrieval days R15-R30 following the final day of training.

#### Behavioral analysis

In all behavioral experiments, we assessed learning and memory recall by calculating the average lick rate difference, which we refer to as the discrimination index (DI). The DI was calculated as follows:

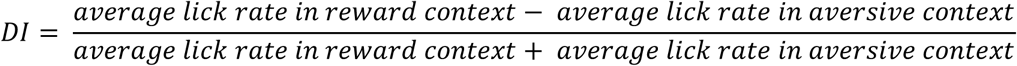

A DI of +1 therefore indicates perfect discrimination in the reward context, while a DI of 0 indicates chance performance. For training sessions, the lick rate was only calculated in the window of 2 seconds preceding the onset of the outcome zone. On retrieval sessions, we included lick rates 1 second before the outcome and during the outcome zone, as no reinforcement is provided but mice still expressed anticipatory licking behavior. Mice used for single cell RNA sequencing, ATAC sequencing, ChIP-sequencing or qPCR were exposed to only the HR or LR condition. We used the area under (AUC) the lick rate curve 2 seconds preceding the start of the outcome zone and up to the end of the outcome zone to assess performance. The AUC was calculated as follows:

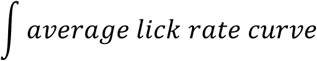

#### Cell culture

The mouse neuroblastoma Neuro2a cell line (ATCC, cat. no. CCL-131) was maintained at 37 °C in a 5% CO_2_ atmosphere. For guide RNA testing, cells were plated at 60% confluency in tissue culture-treated 24 well-plates (Corning, cat. no. 3524) and transfected using Lipofectamine 3000 Transfection Reagent (Thermo Fisher Scientific) following manufacturer’s protocol. Neuro2a cells were cultured in DMEM medium (high glucose, with L-glutamine and sodium pyruvate) containing 10% FBS (Gibco) and 1% Antibiotic-Antimycotic (Gibco).

### CRISPR-Cas9

#### sgRNA design and validation

To design sgRNAs for CRISPR-Cas9 knockout with minimal off-target effects and targeting early exons, 3-8 sgRNAs targeting the protein-coding sequences of selected genes were selected from either CHOPCHOP (http://chopchop.cbu.uib.no) or CRISPick (https://portals.broadinstitute.org/gppx/crispick/public). To select the most efficient sgRNAs per gene, *in vitro* knockout efficiency was assessed by transfecting Neuro2a neuroblastoma cells with pSpCas9(sgRNA)-2A-GFP (Addgene, cat. no. 172221, 1000 ng) using Lipofectamine 3000 Transfection reagent. Neuro2a cells were seeded at 60% confluency in 6-well plate format and transfected over 48 hours, before sorting for eGFP+ cells. DNA was isolated from sorted cells (New England Biolabs, cat. no. T3010) and loci-specific primers were used to amplify loci-specific primers (Supp) to PCR amplify around the edited region with Q5 High-Fidelity 2X Master Mix (New England Biolabs, cat. no. M0492). The DNA band of expected size was agarose gel purified (New England Biolabs, cat. no. T1120) and cleaned up samples were submitted for Sanger sequencing with forward primer (Supp), followed by Synthego ICE Analysis (https://ice.synthego.com/#/) to assess cutting efficiency. DNA amplicons were sequenced using EZ amplicon sequencing (Genewiz) and the percentage of indels calculated. The most efficient guides per gene tested *in vitro* were selected for cloning into AAV:ITR-U6-sgRNA(backbone)-hSyn-Cre-2A-EGFP-KASH-WPRE-shortPA-ITR (pAAV60231) (Addgene, cat. no. 60231). Plasmids to be used *in vivo* were serotyped with AAV9 [coat proteins](https://www.sciencedirect.com/unsupported_browser) and packaged by the University of Arizona Viral Core (at 10E13 GC ml-1 viral titers).

#### sgRNA cloning

For *in vivo* sgRNA design, the sgRNAs were synthesized individually (Integrated DNA Technologies) and cloned into the sgRNA-expressing AAV vector (AAV:ITR-U6-sgRNA(backbone)-hSyn-EGFP-KASH-WPRE-shortPA-ITR, Addgene #60231). Briefly, oligonucleotides (Supp) for each sgRNA were phosphorylated and annealed by T4 Polynucleotide Kinase (New England Biolabs). The sgRNA backbone was digested with SapI (New England Biolabs) and annealed sgRNA inserts were cloned into the backbone by Golden Gate assembly. Then, assembly reactions were transformed into Stable Competent *E. coli* (New England Biolabs, cat. no. C3040H). To verify the sgRNA insert sequences, the sgRNA were Sanger sequenced from the U6 promoter using the U6-fwd primer. For *in vitro* validation and selection of guides, the same steps were carried out, except the oligos were designed with BbsI overhangs and cloned into the sgRNA backbone, pSpCas9(BB)-2A-GFP Px458 (Addgene Plasmid 48138) with BbsI-HF digestion before Golden Gate assembly (New England Biolabs).

#### *In vivo* sgRNA validation

To confirm sgRNA targeting *in vivo*, Rosa26-LSL-Cas9 mice were injected with pAAV60231-sgRNA (Addgene #60231) were housed for three weeks after injection to allow for adequate expression. Then, tissue from ANT or ACC was collected by removing the brain and slicing into 500 µm sections. The brain regions were collected using fluorescence-aided dissection. Tissue sections were either flash frozen in liquid nitrogen or processed immediately. Nuclear isolation proceeded as follows: dissected sections were collected into 500 µl of 54% Percoll buffer. The tissue was homogenized by pipetting with a P1000, followed by a 23G syringe, and a 27G syringe. 5 µl of 10% NP-40 and 5 µl of 10% tween-20 (Millipore Sigma, cat. no. 11332465001) were added to the sample and mixed by pipetting. Samples were placed on ice for 15 minutes, after which 500 µl of 1x buffer, cOmplete EDTA-free protease inhibitor cocktail (Fisher Scientific, cat. no. 11836170001), was added to the tissue. 200 µl of 31% Percoll buffer, followed by 100 µl of 35% Percoll buffer was added to the bottom of the tube, creating a buffer gradient. The samples were centrifuged using a swing rotor for 20,000 xg, 4°C, 10 min. The bottom 100 µl was collected from the bottom layer and blocked by adding 50 µl of 5% BSA/PBS for 5 min on ice. Then, the sample was incubated with primary antibody anti-GFP (BioLegend) at a concentration of 1:2000 for 15 min on ice. The sample was then washed with 1,000 µl of 1x PBS and centrifuged at 500 xg for 5 min and the supernatant was removed. Cells were sorted for GFP+ and DNA from nuclei was isolated (New England Biolabs). To detect percent indel frequency near the targeted stie, primers flanking the deleted region (SX) were used to amplify the intervening region by PCR from genomic DNA, and then Sanger sequenced. Primers were designed to flank a region between 300-500 bp encompassing the sgRNA sequence. Samples were also set for whole-amplicon sequencing (next-generation sequencing) for more detailed mutational characterization.

#### Quantitative PCR (qPCR)

To validate behavioral results of the scRNA pseudotime, an independent behavioral cohort was prepared, and qPCR was performed on tissue dissected from animals for early (R2-R8) or late (R20) time points, exposed to either HR or LR contexts during retrieval (n = 3 - 6 per condition). Animals were sacrificed within 60 min of the retrieval and samples were snap frozen on liquid N_2_. To validate targets of transcriptional regulators, an entirely separate cohort was prepared, where tissue was dissected from control (*Cre*) or knockout animals (*Ash1l* in ACC, *Tcf4* & *Camta1* in ANT). Animals were tested on retrieval on R8 (for *Camta1* and controls), or R20 (*for Tcf4*, *Ash1l* and controls).

Total RNA was extracted from dissected brain tissue using the Total RNA Purification Kit (Norgen Biotek), following the manufacturer’s protocol, with column-based genomic DNA removal to eliminate genomic DNA contamination. RNA concentration and integrity was assessed using an Agilent Bioanalyzer and NanoDrop spectrophotometer. Complementary DNA (cDNA) was synthesized from 500-1,000ng of RNA using the iScript cDNA Synthesis Kit (Bio-Rad) in a 20 μL reaction volume. Primer pairs were designed using NCBI Primer-BLAST to amplify target regions of 80–150 base pairs, with melting temperatures optimized to 60°C ± 1°C and GC content between 40–60%. The specificity of the primers was confirmed by melt curve analysis conducted over a temperature range of 65–95°C with 0.5°C increments. Primer efficiency was validated using a standard curve generated from 10-fold serial dilutions of pooled cDNA, ensuring efficiencies between 90– 110% and an R² value greater than 0.98.

Quantitative PCR reactions were performed in technical triplicates using a QuantStudio 3 Real-Time PCR System, following a 96-well plate format. A master mix was prepared from 2X SYBR Green qPCR Master Mix (Thermo Fisher), nuclease-free H_2_O, and forward and reverse primers, following manufacturer protocol (Table S6). Each final reaction consisted of 17 μL total volume, composed of 15µl of the master mix and 2μL of cDNA template. The cycling conditions included an initial denaturation step at 95°C for 3 minutes, followed by 45 cycles of denaturation at 95°C for 10 seconds and annealing/extension at 60°C for 30 seconds. Melt curve analysis was performed immediately after amplification to confirm primer specificity. Relative quantification of gene expression was calculated using the comparative ΔΔCt method, with normalization to *Map2* as the housekeeping gene, and HR samples normalized to the average of LR samples as a baseline of 1. Genes for which Ct values > 35 were not used. Statistical analyses were performed using GraphPad Prism v9 software. Differences between experimental groups were assessed by a one-tailed, parametric, unpaired t-test, with significance defined as p <0.05. Data are presented as mean ± SEM from 3-6 independent biological replicates (animals).

#### Western Blot

To validate CRISPR knockdown from dissected brain tissue, animals were sacrificed three weeks post-surgery, and the injected region was removed with fluorescence-guided micro-dissection. Brain tissue was lysed using a lysis buffer consisting of 7 mL of RIPA buffer, 70 µL of 100 mM PMSF, and Halt Protease Inhibitor (Thermo Fisher). Tissue samples were placed in a 1.5 mL Eppendorf tube containing 228 µL of lysis buffer and finely minced into small pieces using spring scissors. The minced tissue was homogenized by passing it through a 23G insulin syringe ten times to ensure uniform suspension. An additional 228 µL of lysis buffer and 45 µL of 10% SDS were added to achieve a final SDS concentration of 1%. The mixture was passed through the syringe an additional five times. The homogenized sample was heated at 95°C for 5 minutes to denature proteins, then cooled on ice for 15 minutes. Samples were rotated end-over-end at RT for 40 minutes. Following lysis, samples were centrifuged at 20,000 × g for 20 minutes at 4°C to pellet debris. The supernatant was carefully collected and protein concentration was measured with the Pierce BCA Protein Assay kit and quantified by Nanodrop.

Western blotting was performed using the Odyssey XF Imaging System (LI-COR Biosciences). Cells were lysed in a cold RIPA supplemented with Halt Protease Inhibitor Cocktail (Thermo Fisher), and protein concentrations were determined using a Pierce BCA Protein Assay kit and quantified by Nanodrop. Protein samples (20µg) were denatured by heating at 95°C for 5 minutes in 4X SDS sample buffer with 100 mM DTT and resolved by SDS-PAGE. Proteins were then transferred onto PVDF membranes using a wet transfer system at 70 V for 2 hours. Membranes were blocked in Odyssey Blocking Buffer (PBS-based) for 1 hour at RT. Following blocking, membranes were incubated overnight at 4°C with primary antibodies specific to Tcf4 (Cat no. PA5-88125, Thermo Fisher) diluted 1:1,000 and alpha-Tubulin (cat no. 2144, Cell Signaling Tech) in Odyssey Blocking Buffer containing 0.1% Tween-20. After three washes with PBST, membranes were incubated with IRDye 800CW Goat anti-Rabbit IgG or IRDye 680RD Goat anti-Mouse IgG, diluted 1:15,000 in Odyssey Blocking Buffer + 0.1% Tween-20 for 1 hour at RT in the dark. Membranes were washed three additional times with PBST and imaged while wet on the Odyssey XF Imaging System. Protein bands were visualized in the 700nm and 800nm channels.

#### Nuclear extraction for ATAC-sequencing

Mice were sacrificed within 40-60 minutes from behavioral testing. The anterior thalamus and anterior cingulate of C57Bl6/J mice were dissected, and nuclei were extracted using the percoll gradient method. The dissected tissue was homogenized with 54% percoll (Cytiva) in 1× buffer (50 mM Tris-HCl, pH 7.4; 25 mM KCl; 5 mM MgCl2; 250 mM sucrose) on ice, using a 23 G syringe and a 27 G syringe, 10 strokes each. After homogenization, 10% NP-40 substitute (Roche, cat. no. 11332473001) and 10% Tween-20 were added and incubated for 15 minutes on ice (0.1% final concentration for both NP-40 and Tween-20). Following the incubation, an equal volume of 1× buffer was added and mixed by pipetting. A percoll gradient was prepared by layering 31% and 35% percoll at the bottom of the tube. Nuclei were pelleted in the bottom layer by centrifugation in a swinging bucket (14,000 g, 10 minutes at 4°C). The nuclear pellet was carefully resuspended in blocking buffer (5% BSA/PBS) and incubated for 10 minutes on ice. After blocking, nuclei were labeled by incubation with Alexa Fluor 488-conjugated anti-NeuN antibody (1/2000) for 20 minutes on ice. Nuclei were then washed once in PBS and labeled with 7-AAD (Sigma, cat. no. SML1633). The 7-AAD singlet and NeuN-positive population was sorted by FACS, and the sorted nuclei were used for ATAC-seq.

#### ATAC-sequencing

ATAC-seq preparation was performed as previously described^2^. NeuN-positive nuclei (2-5 × 10^4^) were isolated by FACS and treated with Tn5 transposition mix (Illumina) at 37 ℃ for 30 min in a thermomixer (Eppendorf) at 1,000 rpm mixing. After the Tn5 reaction, DNA was extracted with DNA Clean and Concentrator-5 Kit (Zymo Research, cat. no. D4013) and amplified with NEBNext Ultra II Q5 2× Master Mix (New England BioLabs, cat. no. M0544L). Amplified DNA was extracted using DNA Clean and Concentrator-5 Kit (Zymo Research, cat. no. D4013) and DNA concentration was quantified by NEBNext Library Quant Master (New England BioLabs, cat. no. E7630S). The quality of the purified DNA was checked with a BioAnalyzer (Agilent), and the DNA was sequenced with an Illumina NovaSeq S2 system (50 bp, paired-end).

##### Pre-processing

Paired sequencing reads were 3’ trimmed and filtered for quality (Q≥15) and adapter content using version 0.4.5 of TrimGalore, version 1.15 of cutadapt and version 0.11.5 of FastQC. Version 2.3.4.1 of bowtie2 was used to align reads to mouse assembly mm10 with and duplicates were collapsed to one read using MarkDuplicates in version 2.16.0 of Picard Tools. Enriched regions were discovered using MACS2 with a p-value setting of 0.001, filtered for blacklisted regions (http://mitra.stanford.edu/kundaje/akundaje/release/blacklists/mm10mouse/mm10.blackl ist.bed.gz), and a peak atlas was created using +/-250 bp around peak summits.

##### ATAC-sequencing analysis

Version 1.6.1 of featureCounts was used to build a raw counts matrix and DESeq2 was used to calculate differential enrichment (fold change ≥ 2 and FDR-adjusted p-value ≤ 0.05) for all pairwise contrasts. The BEDTools suite was used to create bigwig files normalized using the DESeq2 sizeFactors, which were also used to normalize the raw counts matrix. Peak-gene associations were created by assigning all intragenic peaks to that gene, while intergenic peaks were assigned using linear genomic distance to transcription start site. Volcano plots were generated using the EnhancedVolcano() package in R with pCutoff = 0.002 and FCcutoff = 1. Network analysis was performed using the assigned genes to differential peaks by running enrichplot::cnetplot in R with default parameters. Motif signatures were obtained using Homer v4.5 on differentially enriched peaks. Heatmaps of ATAC peaks for the transcriptional regulators and their modules were generated using normalized counts filtered by logFold change > 0 from the between R8 vs Homecage pairwise comparison.

### Single-Cell RNA Sequencing

#### Single-Cell Dissociation and Single-Cell RNA Sequencing

Single cell suspensions of anterior prefrontal cortex and anterior thalamus were prepared as described previously^1^. Briefly, mice were sacrificed within 40-60 minutes from behavioral testing, with an overdose of isoflurane, followed by transcardial perfusion with carbonated (95% O_2_, 5% CO_2_) Hanks’ Balanced Salt Solution (HBSS). Brains were removed, 500μm sections were collected, and the region of interest was dissected. The tissue was dissociated using papain (Worthington, cat. no. LS003124) dissolved in Hibernate A buffer (Fisher Scientific, cat. no. NC1787837) and incubated for 25-30 min at 37°C, followed by manual trituration using fire polished Pasteur pipettes and filtering through a 40μm cell strainer (Millipore Sigma, cat. no. BAH136800040). Cells were washed with a wash buffer (PBS + 1% BSA) and centrifuged at 500 g for 5 min, the supernatant was carefully removed, and cells were resuspended in ∼500ul wash buffer and 10% DAPI. Flow cytometry was performed using a BD FACS Aria III Cell Sorter (BD FACSDiva Software, v8.0.1) with a 100-µm nozzle. The cell suspensions were first gated on forward scatter, then within this population based on DAPI-negative cells. Cells were collected in wash buffer, manually counted using a Burker chamber, and suspension volumes were adjusted to a target concentration of 700 -1000 cells/μl. Single cell RNA-sequencing was carried out with the Chromium Next GEM Single Cell 3’ Kit v3.1 (10X Genomics, cat. no. 1000268). Manufacturer’s instructions were followed for downstream cDNA synthesis (12-14 PCR cycles) and library preparation. Final libraries were sequenced on the Illumina NovaSeq S4 platform (R1 – 28 cycles, i7 – 8 cycles, R2 – 90 cycles).

### ChIP-sequencing

#### *In vitro* ChIP antibody validation

Wildtype and knockout Neuro2A cell lines were generated and used to choose the most optimal antibody for ChIP-seq. Briefly, 1×10^^7^ Neuro2A cells were transfected with Px458*-Tcf4-*sgRNA (Addgene #48138) or Px458-*Camta1-*sgRNA (Addgene #48138) for 72 hours and harvested. Freshly harvested cells were fixed with 1% formaldehyde for 10 min, after which the reaction was quenched by the addition of glycine to the final concentration of 0.125 M. Fixed cells were washed twice with PBS, snap frozen and store at -80°C.

#### Behavior for ChIP-seq

For *in vivo* ChIP-experiments, an independent behavioral cohort was run according to the behavioral task above. Cohorts of Rosa26-LSL-Cas9 animals were injected with either *Ash1l-*sgRNA, *Tcf4-*sgRNA, or *Camta1-*sgRNA. For assessing the histone methylation landscape over time, both control animals and *Ash1l-*KO animals were tested on R2, R8, and R20. For *Camta1-*KO and *Tcf4-*KO, animals were tested and sacrificed on R8. For *Ash1l-*KO and controls used for H3K4me3 ChIP, two independent biological replicates were included, each pooled from tissue dissected from 3-4 animals. For *Camta1-*KO and *Tcf4-*KO (and corresponding R8 controls), one biological replicate was included, each pooled from 9 mice to obtain enough cells for downstream ChIP-seq for those specific antibodies, as determined by the *in vitro* ChIP input.

#### Mouse brain nuclei isolation of neurons for ChIP-seq

Animals were anesthetized with isoflurane within an hour of performing the task retrieval and perfused with PBS. ANT or ACC tissue was micro-dissected under a microscope and snap frozen on liquid nitrogen for later processing. Nuclei were isolated from mouse brain tissue based on a previously described method^18^ with slight modifications. Tissue from 8-9 mice were pooled for Camta1-knockout and Tcf4-knockout, and their respective controls, and tissue from 2-3 mice were pooled for each biological replicate for *Ash1l-* knockout testing. Briefly, 0.3 ml fixative solution (1% (wt/vol) formaldehyde in DPBS) was added to the pooled sample (∼60-80mg tissue) in a 1.5 ml tube. Tissue was gently homogenized in the fixative by gently breaking up with a P1000 pipette tip until no large visible pieces remained. For Ash1l samples, the 0.3 ml homogenate was added to a 4.7 ml fixative solution in a 15 ml tube using a P1000 tip, washing out the 1.5 ml tube to maximize homogenate transferred. Homogenate was rocked at 30 cycles/min, RT for 10 min. To quench the formaldehyde and prevent overfixation of the sample, 0.5 ml 2.5 M glycine, [final] = 0.125 M, was added to the homogenate and rocked at 30 cycles/min, RT for 5 min. (For Tcf4 and Camta1 samples, a double fixation protocol was used: homogenization of sample in 0.3 ml 2 mM DSG solution, followed by transfer to 4.7 ml 2 mM DSG, rocked at 30 cycles/minute, RT, 30 min. 37% (wt/vol) formaldehyde was added to the homogenate to create a 1% (wt/vol) formaldehyde solution, rocked for 10 min at RT. The sample was quenched with 0.125 glycine.) After fixation and quenching, the rest of the steps were the same. The homogenate was spun at 1,100*g* for 5 min, 4°C in a swing rotor bucket centrifuge with 15 ml adaptors. Supernatant was removed and pellet was suspended with 10 ml ice-cold NF1 buffer (10 mM Tris-HCl pH 8.0, 1 mM EDTA, 5 mM MgCl_2_, 0.1 M sucrose, and 0.5% (vol/vol) Triton X-100) for a first wash. This was repeated for a second wash and on the third (final) was, sample was re-suspended in 5 ml ice cold NF1 buffer. Homogenate was incubated on ice for 30 min, then transferred to a 7 ml glass dounce. A loose pestle was passed through the homogenate 20x, and then a tight pestle passed through 5x. The homogenate was passed through a 70µm strainer and collected into a 50 ml tube. Ann additional 15 ml NF1 buffer was passed through the strainer to collect a total volume of 20 ml. A 5 ml sucrose cushion (1.0 M sucrose cushion with 1.2 M sucrose solution with 10 mM Tris-HCl pH 8.0, 3 mM MgCl_2_, and 1 mM DTT) was laid underneath the homogenate by slowly pipetting to the bottom of the tube with a P1000, with care to maintain an interphase. The sucrose gradient was spun at 3,900*g* for 30 min at 4°C. After spinning, a white nuclei pellet was visible at the bottom, below the sucrose cushion. The upper, aqueous phase, interphase, and sucrose cushion were gently removed. The nuclei pellet was resuspended in 1 ml NF1 buffer and transferred to a 15 ml tube. 9 ml NF1 buffer was added to wash the nuclei pellet, followed by spinning at 1,600*g* for 5 min 4°C to pellet the nuclei. For antibody staining for nuclei sorting, the nuclei pellet was resuspended in 5 ml FANS buffer, spun at 1,600 g for 5 min, 4°C, and resuspended in 0.3 ml FANS buffer. 3µl was taken for an unstained control and the remaining nuclei were transferred to a FACS tube for staining overnight 4°C with NeuN-Ax488 antibody (Millipore, #MAB377X) at 1:2500 dilution with gentle rocking. The next day, 4 ml FANS buffer was added to the nuclei, then spun at 1,600*g* for 5 min, 4°C. The remaining 0.3 ml nuclei were passed through a 35 µm cap into a new tube for FACS, and counterstained with 0.5 µg ml^−1^ DAPI. Nuclei were sorted using a 100 µm nozzle. Nuclei were gated based on DAPI+, NeuN+ according to the original protocol. After sorting, nuclei samples were adjusted to 1% wt/vol BSA, then spun at 1,600*g* for 15 min 4°C to collect nuclei for further processing. FANS buffer was removed and samples were snap frozen on liquid N_2_.

#### Chromatin immunoprecipitation (ChIP) and sequencing

Frozen pellets of fixed cells (*in vitro*) or fixed, dissociated nuclei (*in vivo* behavior) were resuspended in SDS buffer (100 mM NaCl, 50 mM Tris-HCl pH 8.0, 5 mM EDTA, 0.5% SDS, 1x protease inhibitor cocktail from Roche). The resulting nuclei were spun down, resuspended in the immunoprecipitation buffer at 1 mL per 0.5 million cells (SDS buffer and Triton Dilution buffer (100 mM NaCl, 100 mM TrisHCl pH 8.0, 5 mM EDTA, 5% Triton X-100) mixed in 2:1 ratio with the addition of 1x protease inhibitor cocktail (Millipore Sigma, #11836170001) and processed on a Covaris LE220+ Focused-ultrasonicator to achieve an average fragment length of 200-300 bps with the following parameters: PIP=420, Duty Factor=30, CBP/Burst per sec=200, Time = 20min. Chromatin concentrations were estimated using the Pierce™ BCA Protein Assay Kit (ThermoFisher Scientific #23227) according to the manufacturer’s instructions. The immunoprecipitation reactions were set up in 500uL of the immunoprecipitation buffer in Protein LoBind tubes (Eppendorf, #22431081) and pre-cleared with 50uL of Protein G Dynabeads (ThermoFisher Scientific #10004D) for two hours at 4°C. After pre-clearing, the samples were transferred into new Protein LoBind tubes and incubated overnight at 4°C with 5ug of TCF4 (ProteinTech, #22337-1-AP), CAMTA1 (Millipore, #SAB4301068) or H3K4me3 antibody (Epicypher, 13-0041 or 13-0060)]. The next day, 50uL of BSA-blocked Protein G Dynabeads were added to the reactions and incubated for 2 hours at 4°C. The beads were then washed two times with low-salt washing buffer (150mM NaCl, 1% Triton X-100, 0.1% SDS, 2mM EDTA, 20mM TrisHCl pH8.0), two times with high-salt washing buffer (500mM NaCl, 1% Triton X-100, 0.1% SDS, 2mM EDTA, 20mM TrisHCl pH8.0), two times with LiCL wash buffer (250mM LiCl, 10mM TrisHCl pH8.0, 1mM EDTA, 1% Na-Deoxycholate, 1% IGEPAL CA-630) and one time with TE buffer (10mM TrisHCl pH8.0, 1mM EDTA). The samples were then reverse-crosslinked overnight in the elution buffer (1% SDS, 0.1 M NaHCO_3_) and purified using the ChIP DNA Clean & Concentrator kit (Zymo Research, #D5205) following the manufacturer’s instructions. After quantification of the recovered DNA fragments, libraries were prepared using the ThruPLEX®DNA-Seq kit (R400676, Takara) following the manufacturer’s instructions, purified with SPRIselect magnetic beads (B23318, Beckman Coulter), and quantified using a Qubit Flex fluorometer (ThermoFisher Scientific) and profiled with a TapeStation (Agilent). The libraries were sent to MSKCC Integrated Genomics Operation core facility for sequencing on an Illumina NovaSeq 6000 (aiming for 30-40 million 100bp paired-end reads per library).

#### ChIP-sequencing data processing and analysis

Paired sequencing reads were 3’ trimmed and filtered for quality (Q≥15) and adapter content using version 0.4.5 of TrimGalore (https://www.bioinformatics.babraham.ac.uk/projects/trim_galore) and running version 1.15 of cutadapt and version 0.11.5 of FastQC. Version 2.3.4.1 of bowtie2 (http://bowtie-bio.sourceforge.net/bowtie2/index.shtml) was used to align reads to mouse assembly mm10 with and duplicates were collapsed to one read using MarkDuplicates in version 2.16.0 of Picard Tools. Enriched regions were discovered using MACS2 (https://github.com/taoliu/MACS) with a p-value setting of 0.001, filtered for blacklisted regions (http://mitra.stanford.edu/kundaje/akundaje/release/blacklists/ mm10-mouse/mm10.blacklist.bed.gz), and a peak atlas was created using +/-250 bp around peak summits for ATAC data and the entire enriched region for ChIP data. Version 1.6.1 of featureCounts (http://subread.sourceforge.net) was used to build a raw counts matrix and DESeq2 was used to calculate differential enrichment (fold change ≥ 2 and FDR-adjusted p-value ≤ 0.05) for all pairwise contrasts. The BEDTools suite (http://bedtools.readthedocs.io) was used to create ATAC bigwig files normalized using the DESeq2 sizeFactors, which were also used to normalize the raw counts matrix. For histone modification data, the ChIP signal was normalized to the sequencing depth for uniquely mapped reads, while TF ChIP data were normalized to an external spike in by scaling the data inversely to the number of Drosophila H2Av reads. Peak-gene associations were created by assigning all intragenic peaks to that gene, while intergenic peaks were assigned using linear genomic distance to transcription start site.Network analysis was performed using the assigned genes to differential peaks by running enrichplot::cnetplot in R with default parameters. Gene set enrichment analysis (GSEA, http://software.broadinstitute.org/gsea) was performed with the pre-ranked option and default parameters by assigning a gene to the single peak with the largest (in magnitude) log2 fold change associated with it for each paired contrast. Composite and tornado plots were created using deepTools v3.3.0 by running computeMatrix and plotHeatmap on normalized bigwigs with average signal sampled in 25 bp windows and flanking region defined by the surrounding 2 kb. Motif signatures were obtained using Homer v4.5 (http://homer.ucsd.edu) on differentially enriched peaks.

### Photometry

#### Data Acquisition & Post processing

A custom multi-fiber photometry setup, as previously described^31^, was used to simultaneously record bulk calcium signals from ANT and ACC while mice performed the VR-based contextual discrimination task. We recorded at 11 Hz with an excitation wavelength of 470 nm. Prior to each session, mice were head fixed, and each optical cannula was cleaned with 70% ethanol. For analysis, the images captured were post-processed using custom MATLAB scripts. Regions of interest were manually drawn for each fiber to extract fluorescence values throughout the experiment. Raw signals were high-pass filtered and then z-scored.

#### Data analysis

All subsequent photometry data analysis was carried out using custom Python scripts. To calculate Pearson correlations between ANT and ACC, signals from the cue and reinforcement zone were concatenated. To focus on correlations between relevant calcium events rather than noise, only signals above a 0.5 magnitude were used. Pearson correlations were then calculated using the Scipy *pearsonr* function. To estimate mutual information between ANT and ACC within each session, the Scikit-learn function *mutual_info_regression* was used. Finally, to assess the complexity of local ANT and ACC calcium signals across experimental groups, sample entropy was calculated using the AntroPy *sample_entropy* function with embedding dimension (m) = 2 and tolerance (r) = 0.2

#### Single-Cell RNA Sequencing Data Analysis

##### QC and visualization

RAW sequencing reads were aligned to the GRCm38/mm10 mouse reference genome and all scRNA-seq datasets were initially processed individually using the Sequence Quality Control (SEQC) package^4^. The SEQC pipeline performs cell barcode and UMI correction to generate a count matrix (cells x genes); true cells were distinguished from empty droplets based on the cumulative distribution of total molecule counts and cells with a high fraction of mitochondrial molecules were filtered (> 20%), as these are likely apoptotic. The Python Scanpy package (v1.9.3) was used to further analyze the data^5^. Presence of doublets was verified again using Scrublet^6^ and cell barcodes from Cellplex (10x Genomics, cat. no. 1000261) were identified for each biological replicate. After this initial preprocessing, samples from all time points were merged. Cells with <1,500 UMIs per cell and <1,000 genes per cell, and genes detected in < 10 cells were removed. Standard median library size normalization followed by log transformation (pseudo-count = 1) was applied on the data. Ribosomal and mitochondrial genes were removed. Next, highly variable gene (HVG) selection was performed using Scanpy’s scanpy.pp.highly_variable_genes() function with the seurat_v3 method on raw gene expression counts, and Principal Component Analysis was applied to reduce the dimensionality to 30 principal components to obtain 4000 highly variable genes (HVGs). A nearest neighbor graph (n_neighbors = 30) was computed between cells using these principal components, and PhenoGraph Leiden^7^ clustering was applied on the PC space (with *k* = 30). We established that clustering was robust to slight changes in *k* by re-clustering the cells under varying *k* and measuring consistency using the adjusted Rand index (sklearn package in Python). Datasets were visualized with UMAP embeddings computed on the obtained principal component analysis (PCA).

##### Cell type assignment

To assign cell type labels, we manually assessed patterns of mean marker genes expression across clusters using custom marker genes^3^ (**Table S1,** extended data fig. 2e, 2g). We also calculated differentially expressed genes (DEGs) for each cluster versus all other clusters with Scanpy’s scanpy.tl.rank_gene_groups() function using the Wilcoxon rank sum test with Benjamini-Hochberg correction to identify top expressed genes in clusters that were not easily identifiable. Once the neuronal cluster was identified, it was subsetted and re-clustered using the first 30 principal components restricted to 2100 HVGs. A set of canonical marker genes for anterior/posterior thalamic nuclei were used in the ANT samples and another set of marker genes to identify cortical layers was used in the ACC samples^3,8^ (**Table S2**).

##### Differential gene expression and Gene ontology pathway analysis

Differential gene expression analysis between the HR and LR condition across time points was performed using the “MAST” R package^9^ on log-normalized values, and DEGs were defined as having an adjusted p value < 0.05 and log fold change > 0.2. Gene set enrichment analysis was performed using the fast gene set enrichment analysis (fGSEA) package (fgsea v1.18.0), and the GO_Biological_Function_2023 gene set library was used for gene ontology analysis^10^.

##### Wasserstein distance computation

Global transcriptional distances between HR and LR samples across consecutive time points were estimated by calculating the Wasserstein distances using the Pertpy package^11^, and the pt.tl.Distance(“wasserstein”) function using obsm_key=“X_pca”. Transcriptional distances restricted to the DEGs were estimated using custom code based on the “Pertpy” algorithm.

##### Pseudo-time analysis and visualization

To infer changes in cell states in the neuronal population from early learning state throughout consolidation, we subsetted the ANT and ACC neurons and included clusters 0, 1,4,2, 5 for ANT and clusters 0, 1, 2, 3, 7 for ACC, as these clusters were the closest to each other on the UMAP space and likely represent more phenotypically similar states. We recalculated highly variable genes using the scanpy.pp.highly_variable_genes() function with the seurat_v3 method to obtain 1000 HVGs. A nearest neighbor graph (n_neighbors = 30) was computed between cells using 30 principal components. We then applied pseudo-time trajectory estimation. We chose to use 1) the Palantir algorithm which assumes a known starting point and a unidirectional progression from a “less- to a more-differentiated state” hence orders cells along a continuum trajectory and assigns each cell a probability for reaching each terminal state^12^. We then computed diffusion maps with Palantir (n_components=5, n_waypoints = 500) to identify major axes of variation. Once the diffusion maps were computed, Palantir computes rescaled diffusion components on which the trajectory inference is performed. We selected a random starting cell from “Training day 2” for the ANT sample and from “Test day 2 HR” for the ACC sample. We also ran another trajectory analysis, 2) CellRank with the transition matrix computed using the Cytrotrace implementation^13^. Cytotrace assumes the number of genes expressed per cell as a signal of “differentiation”. We observed that with both algorithms the “terminal states’’ were not the latest sampled time points suggesting that the extreme branches represented extreme phenotypes rather than terminal states.

In addition to computing pseudo-time, Palantir also visualizes the data using tSNE on the multiscale diffusion components (Fig. 3a, 3e). We used this visualization to study shifts in population density along time points per condition (HR or LR). The density plots were made using the Python’s Seaborn package and kdeplot() function. To estimate percent of densities for neurons from each condition and time, we calculated the density of neurons in an embedding per condition using Scanpy’s scanpy.tl.embedding_density() and visualized using scanpy.pl.embedding_density(). In a similar way, densities of Fos+ neurons were also plotted on the trajectory tSNE embedding to observe shifts between samples. A neuron was defined as being Fos+ if the expression of Fos gene was greater than 0.25. To visualize genes correlated with the “early” and “late” consolidation branches and hence estimate fate probabilities, we used CellRank with the Palantir implementation and trajectory obtained previously with Palantir, using cr.kernels.PseudotimeKernel() with time_key=’pseudotime_palantir’.

##### MiloR analysis

To quantify densities of HR vs LR neurons along the pseudo-time trajectory and identify statistically significant enrichments, we used the MiloR^15^ algorithm which groups cells into partially overlapping local neighborhoods and computes differential neighborhood abundances across conditions (here HR versus LR). We constructed a k Nearest Neighbor (kNN) graph (*k* = 30) on principal components using the buildGraph() function and then constructed neighborhoods on the kNN graph using the makeNhoods() function with default parameters (prop = 0.1, refined = true). The number of neurons present in each neighborhood were quantified using the countCells() function and the statistical significance was assessed using testNhoods() and calcNhoodDistance() for spatial FDR correction. We visualized the distribution of log fold changes in each condition using the plotNhoodGraphDA() function with alpha set to 1 in all cases.

##### Gene target module estimation

To estimate the putative target genes of the identified transcriptional regulators, we used the Chea3 browser^16^, which assembles TF-target gene set libraries, and in this case, we used the Enrichr library. For each time point, we inputted the list of DEGs for the HR condition and identified the same transcriptional regulators derived from the pseudo-time lineage correlation analysis. From the analysis, we obtained as list of overlapping genes predicting each transcriptional regulator.

##### Comparison of expression of TRs across datasets

To compare expression levels of the identified ANT and ACC TRs, we downloaded the corresponding prefrontal cortex and thalamus datasets from the Mouse Brain Atlas (MBA) from the Linnarsson lab^3^ as well as from the Allen Brain Atlas^17^ (ABC). Pre-processing was carried out for the MBA dataset in a similar manner as described earlier, and the ABC dataset version 10xv2-log2 version was retrieved. Both datasets were clustered using Phenograph, and violin plots were obtained using the scanpy.pl.violin() function with default parameters.

### Normality tests

To assess the distribution of our data, we performed normality and lognormality tests using GraphPad Prism (v9.5.1). For each dataset, normal quantile-quantile (QQ) plots were generated to visually evaluate the fit to a Gaussian or lognormal distribution. Formal statistical tests for normality (Shapiro-Wilk) were also applied, as implemented in Prism. QQ plots were used alongside p-values from these tests to guide the choice of appropriate statistical analyses. Data failing both normal and lognormal tests were analyzed using non-parametric methods.

---

**Extended data Fig. 1:**
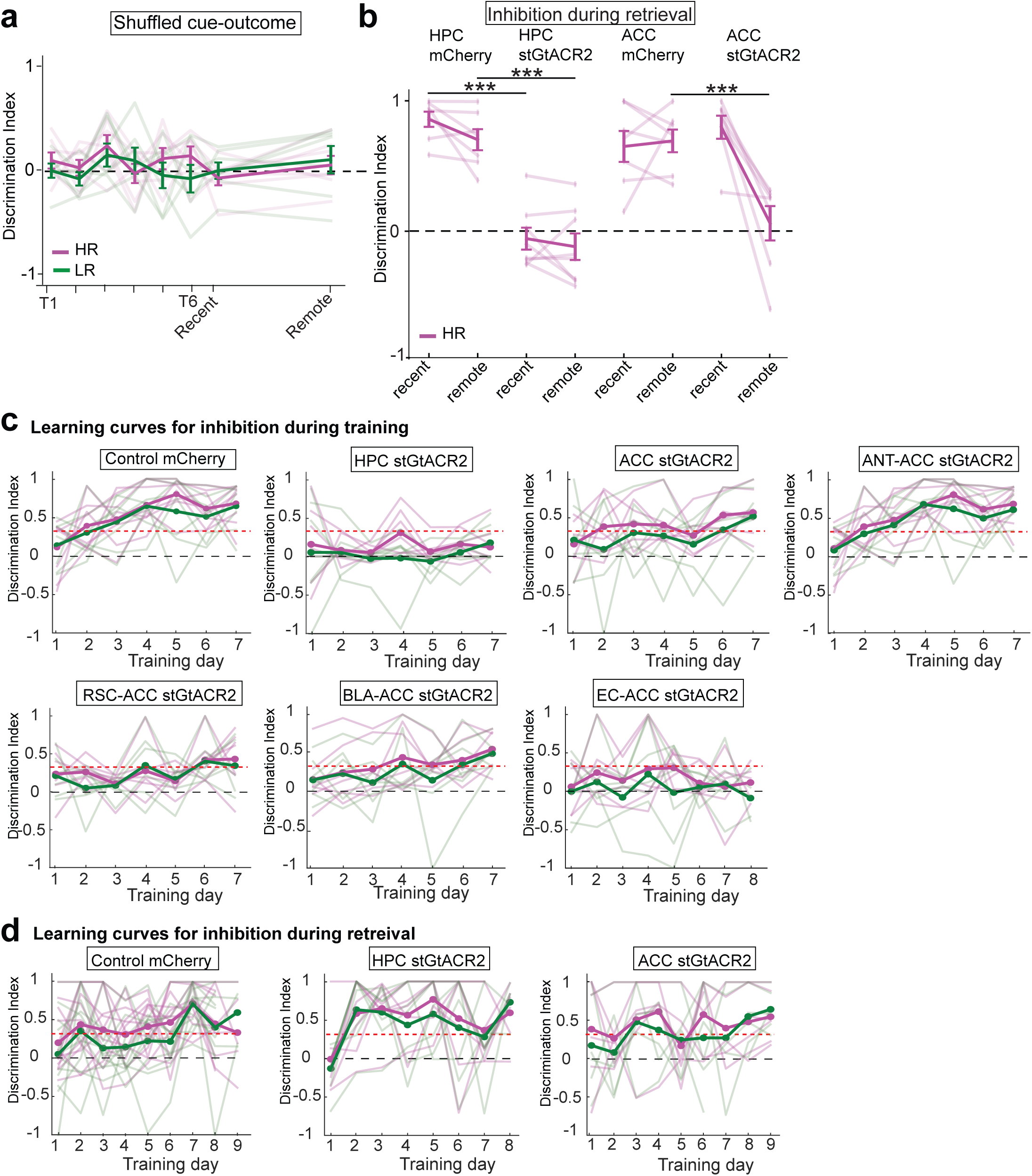
Learning and retrieval performances during shuffled cue-outcome zones, inhibition during training and retrieval. **a**, Discrimination indices for learning and retrieval performance of mice exposed to shuffled cue-outcomes, *n* = 7 mice, individual data points shown (faded lines), with mean ± SEM (solid line). **b**, Optogenetic inhibition during recent and remote retrieval, in control mCherry, HPC stGtACR2 and ACC stGtACR2. Light delivered during cue periods of each trial. Quantification of discrimination indices between HR and aversive lick rates, *n* = 7-8 mice per cohort, individual data points shown, *** *P*<0.0001 between HPC mcherry and HPC stGtACR2 on recent and remote retrieval, \*\*\**P* = 0.0001 between ACC mcherry and ACC stGtACR2 on remote retrieval, One-way ANOVA with Bonferroni correction. **c**, Discrimination indices for learning performance in HR or LR for mice receiving inhibition during training days, *n* = 7-9 mice per cohort, individual data points shown (faded lines), with mean ± SEM (solid line). **d**, Discrimination indices for learning performance in HR or LR for mice receiving inhibition during retrieval days, *n* = 7-8 mice per cohort, individual data points shown (faded lines), with mean ± SEM (solid line).

**Extended data Fig. 2:**
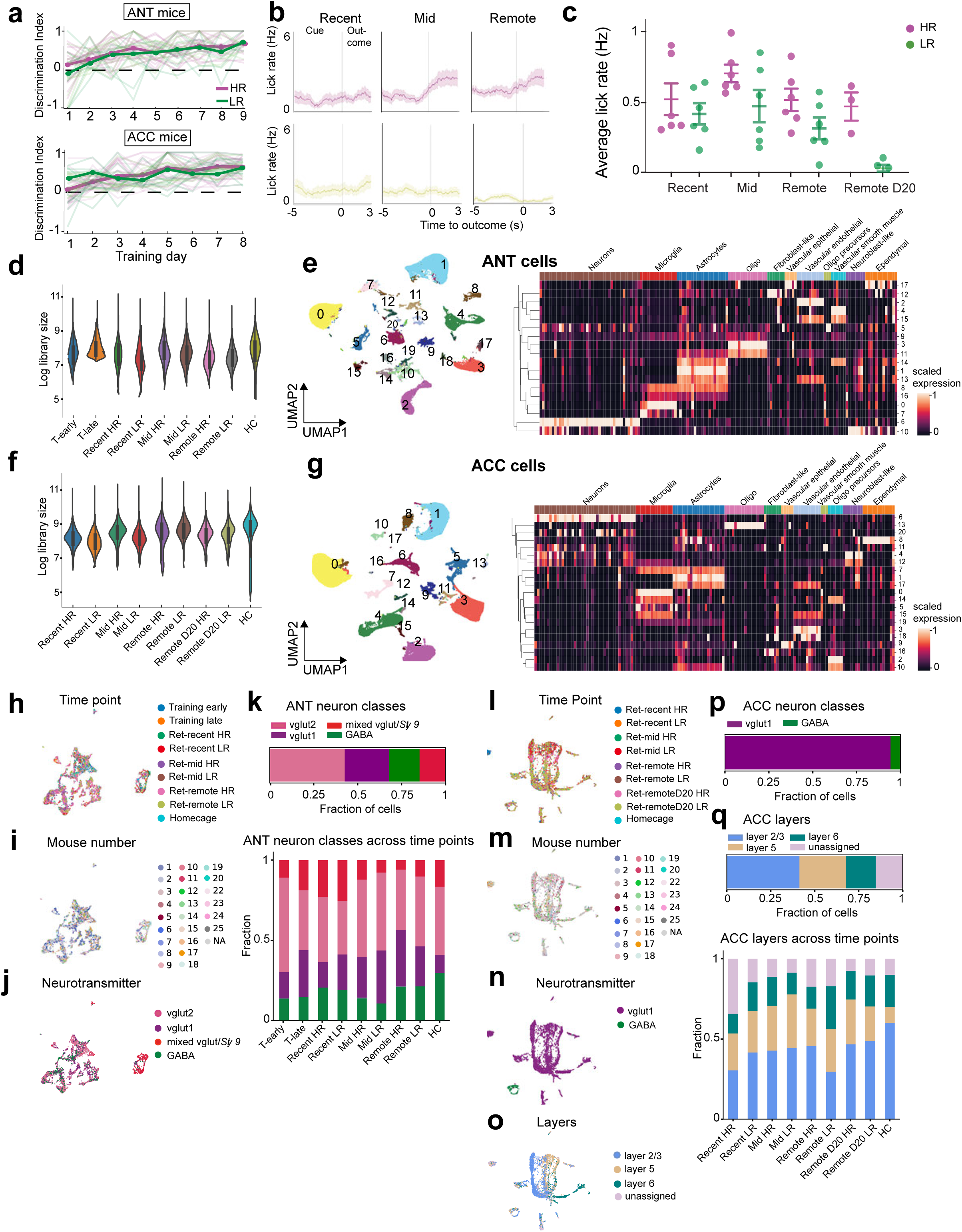
scRNA-sequencing behavioral data: cell typing and sub-setting of neurons. **a**, Discrimination indices for learning performance in HR and LR contexts of mice used for scRNA sequencing, *n* = 23 mice for ANT and 24 mice for ACC, individual data points shown (faded lines), with mean (solid line). **b**, Representative lick traces from one mouse showing trial averages in HR and LR on recent, mid and remote retrieval, data are mean (solid line) ± SEM (shaded area), *n* = 30-40 trials. **c**, Memory performance in HR and LR contexts for mice used for scRNA sequencing, \**P =* 0.027 One-way ANOVA with Bonferroni correction. **d, f**, Library size for ANT (**d**) or ACC (**f**) samples. **e, g**, Left: UMAP visualization of all cells from ANT (*n* = 176566 cells) or ACC (*n* = 145327 cells), clustered based on transcriptional profile and colored by cluster number. Right: heatmap of expression of canonical marker genes for cell types across clusters. **h, l**, UMAP sub-clustering of cells identified as neurons in ANT (*n* = 5535 cells) (**h**) or ACC, (*n* = 5671 cells) (**l**) colored by time point. **i**, **m**, UMAP of ANT (**i**) or ACC (**m**) neurons colored by feature barcode of biological replicates. **j**, **n**, UMAP of ANT (**j**) or ACC (**n**) neurons colored by neurotransmitter class. **k**, Top: proportion of ANT neurons assigned to each neurotransmitter class. Bottom: breakdown of neuron classes across time points and conditions. **o**, UMAP of ACC neurons colored by cortical layer assignment. **p**, Proportion of ACC neurons assigned to each neurotransmitter class. **q**, Top: proportion of ACC neurons assigned to each cortical layer. Bottom: breakdown of cortical layer assignment across time points and conditions.

**Extended data Fig. 3:**
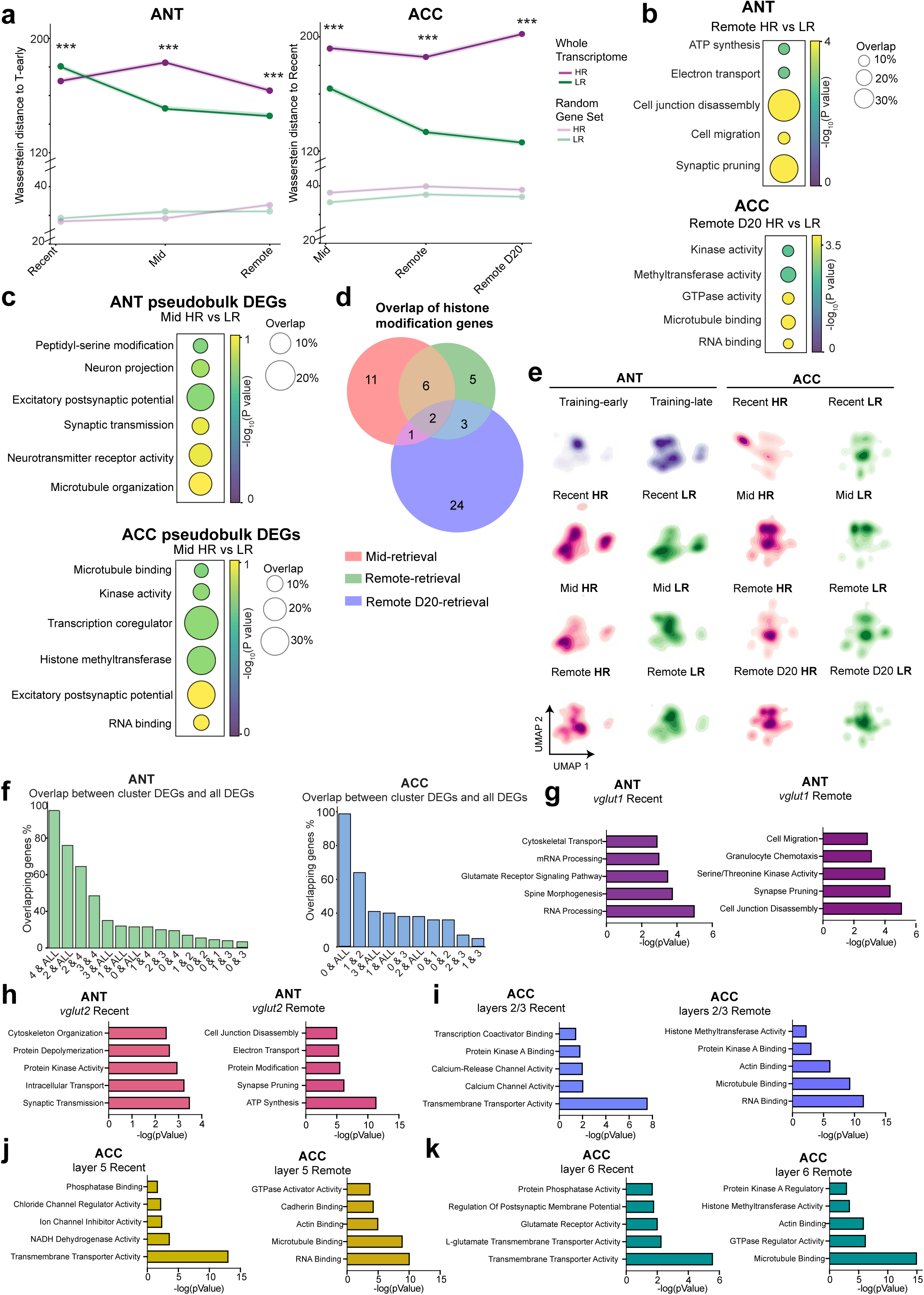
ANT and ACC recruit distinct gene programs during memory persistence. **a**, Line plots of Wasserstein global transcriptional distance, using the whole transcriptome (solid colors) or a random set of genes (faded). Distances are to early-training in ANT (left) and to recent-retrieval in ACC (right), \*\*\**P* < 0.0001 for global distances for recent, mid and remote HR vs LR in ANT, \*\*\**P* < 0.0001 for global distances for mid, remote and remote D20 HR vs LR in ACC, One-way ANOVA with Bonferroni correction. **b**, Gene ontology (GO) analysis of genes upregulated in HR neurons on remote retrieval in ANT (top) and remote retrieval day R20 in ACC (bottom). **c**, GO analysis of DEGs obtained using pseudobulk differential analysis. **d**, Overlap of genes belonging to the histone methylation module across retrieval time points in ACC. **e**, Density plots showing distribution of neurons per conditions across UMAP clusters in ANT (left) or ACC (right). **f**, Proportion of overlap between mid-retrieval DEGs using all cells from each condition and mid-retrieval HR vs LR DEGs using cells from each cluster. Comparisons are between clusters (numbered), or between per-cluster DEGs and all-cells DEGs (ALL). Shown for ANT (left) and ACC (right). **g**, **h**, GO analysis of DEGs from ANT neurons classified as *vglut1* (**g**) or *vglut2* (**h**). **i**, **j**, **k**, GO analysis of DEGs from ACC neurons classified as *layer 2/3* (**i**), *layer 5* (**j**) or *layer 6* (**k**).

**Extended data Fig. 4:**
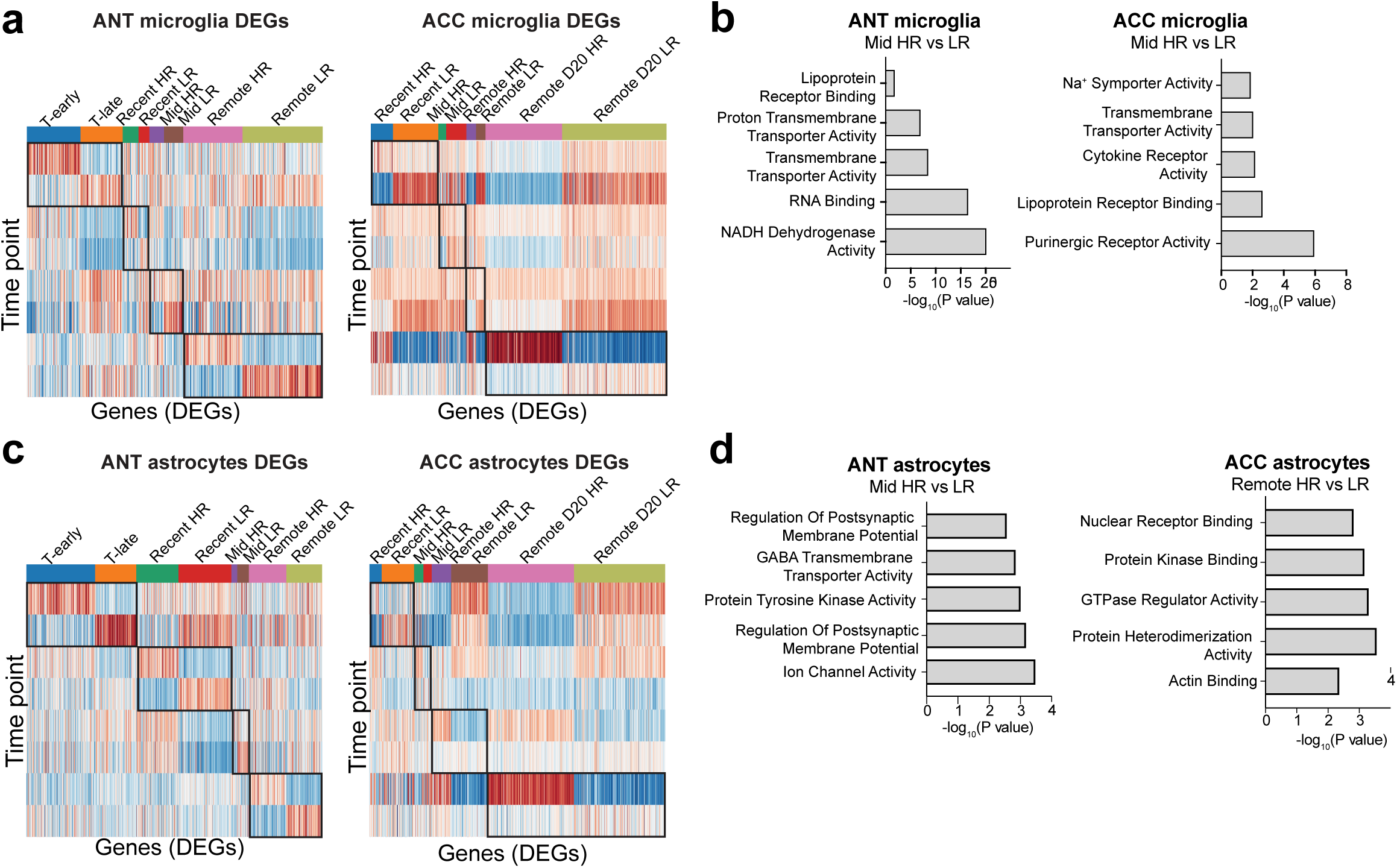
Transcriptional changes in microglia and astrocytes. **a**, Heatmap of z-scored expression of DEGs in each condition in ANT microglia (left) or ACC microglia (right) across time points. Columns are DEGs, rows are time points, *n* = 26-423 DEGs per time point. **b**, GO analysis of mid-retrieval HR vs LR DEGs from ANT (left) or ACC (right) microglia. **c**, Heatmap of z-scored expression of DEGs in each condition in ANT astrocytes (left) or ACC astrocytes (right) across time points. Columns are DEGs, rows are time points, *n* = 24-270 DEGs per time point. **d**, GO analysis of mid and remote retrieval HR vs LR DEGs from ANT (left) or ACC (right) astrocytes.

**Extended data Fig. 5:**
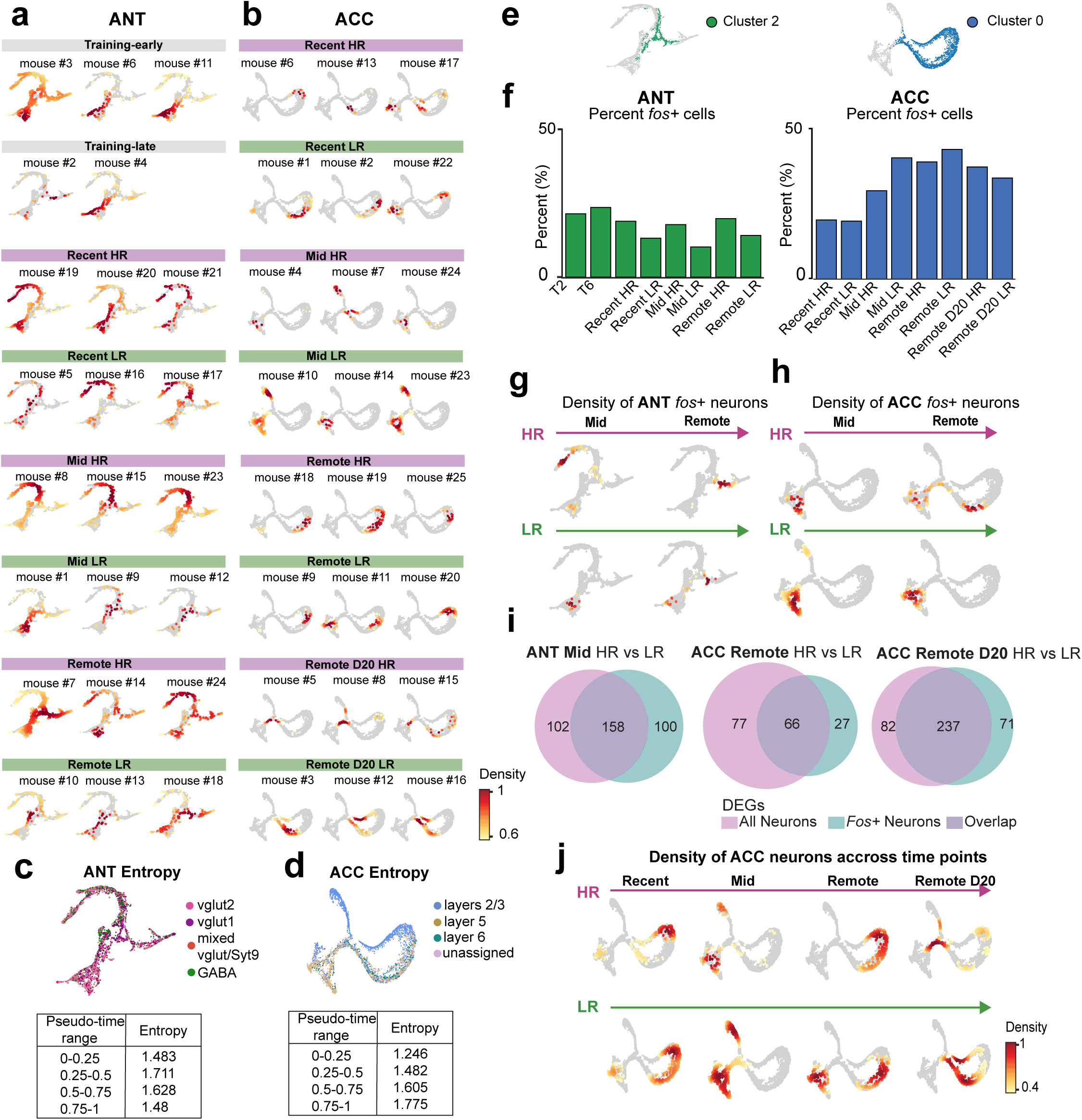
Pseudo-time trajectories capture macrostates associated with memory persistence. **a, b,** Kernel density estimate tSNE plots of ANT (**a**) or ACC (**b**) neurons for each biological replicate (*n* = 2-3 mice per time point) across all time points. **c**, **d**, Top: tSNE visualization of ANT (**c**) or ACC (**d**) pseudo-time trajectory colored by neuronal class in ANT and anatomical layer in ACC. Bottom: Entropy values reflecting heterogeneity of cell classes or layer assignment along the pseudo-time trajectory. **e,** tSNE visualization of ANT (left) or ACC (right) pseudo-time trajectories highlighting the cluster with greatest DEGs overlap with all mid-retrieval DEGs. **f**, Bar plot of percent *Fos*+ neurons across time points in ANT (left) or ACC (right) neurons used to construct pseudo-time trajectories. **g**, **h**, tSNE visualization of ANT (**g**) or ACC (**h**) trajectories, colored by *Fos*+ neurons with scored density > 0.6. **i**, Representative Venn diagrams of shared genes between DEGs obtained using all cells per condition and DEGs obtained using *Fos*+ cells per condition. Shown for mid—retrieval in ANT (left) and mid and remote retrieval in ACC (right). **j**, Density plots for HR and LR ACC neurons on the tSNE pseudo-time space across retrieval days.

**Extended data Fig. 6:**
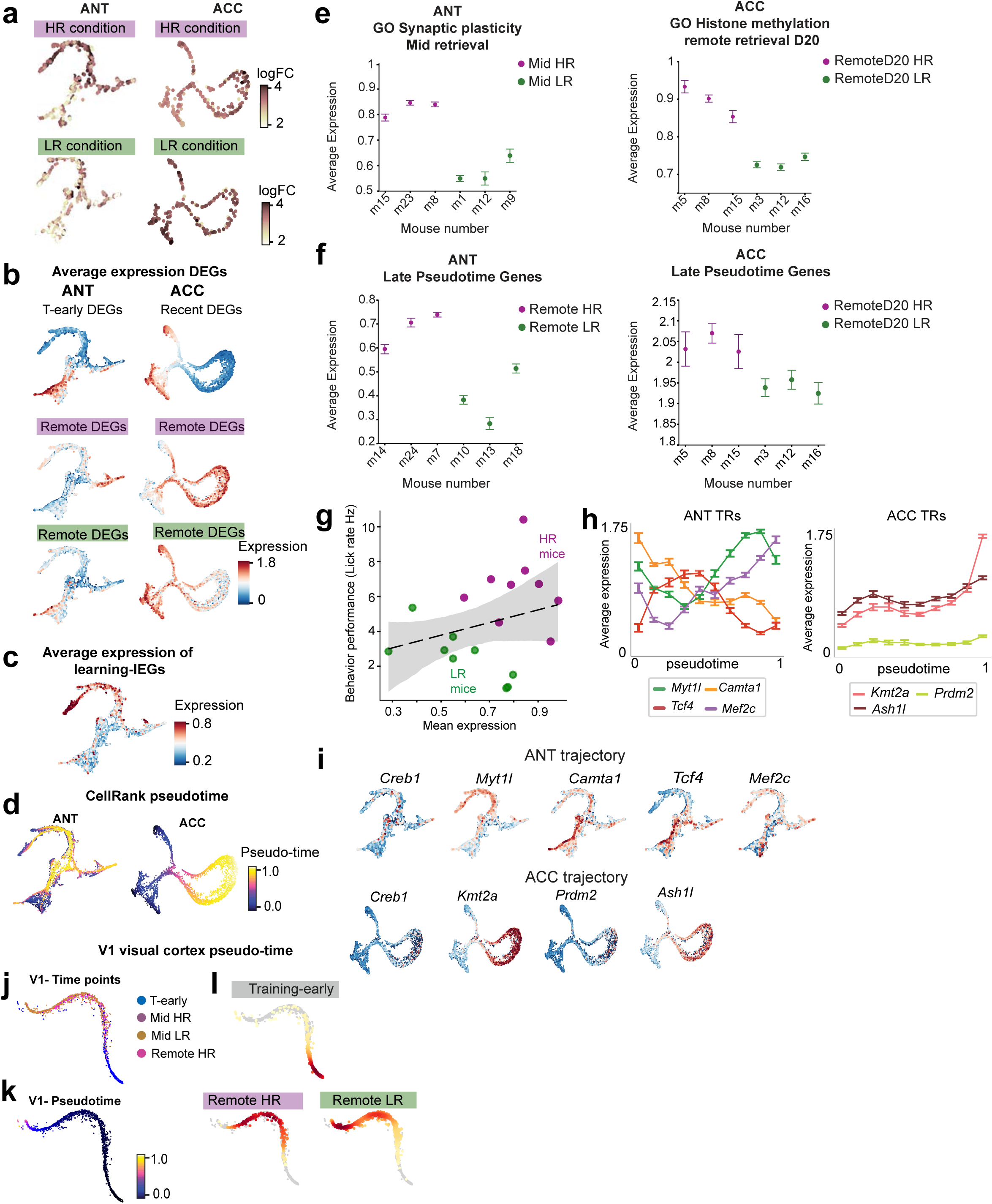
Expression of DEGs, GO modules and macrostate correlated genes along pseudo-time trajectories. **a**, tSNE visualizations of ANT (left) and ACC (right) neurons, colored by abundance of cells from HR or LR conditions. **b,** tSNE visualizations of ANT (left) and ACC (right) neurons, colored by average expression of DEGs from early-training and remote retrieval for ANT or recent-retrieval and remote retrieval day 20 for ACC (HR in magenta and Lr in green, units log_2_ CPM +1). **c**, tSNE visualization of ANT neurons colored by average expression of adhesion or structural related genes (units log_2_ CPM +1). **d**, tSNE representations of pseudo-time trajectories for ANT (left) or ACC (right) neurons using the CellRank algorithm. Stars highlight apex points detected by CellRank. **e**, Expression of genes associated with the synaptic plasticity GO module in ANT (left) and the histone methylation GO module in ACC (right) across HR and LR biological replicates. Sown for mid-retrieval in ANT and remote retrieval DR20 in ACC (units log_2_ CPM +1). **f**, Expression of genes correlated with the early in ANT (left) or late in ACC (right) trajectory macrostates across HR and LR biological replicates. **g**, Pearson correlation between behavioral performance during retrieval (measure as lick rate Hz) and average expression of genes shown in (**e**) and (**f**), each dot is a mouse, mice tested on the HR condition in magenta and mice tested on the LR condition in green, r = 0.27. **h,** Average expression of ANT or ACC transcriptional regulators in ANT neurons (left) or ACC neurons (right) along the ANT or ACC pseudo-time trajectory, respectively. Data are mean (solid line) ± SEM (error bars). **i**, Average expression of ANT (top) or ACC (bottom) TRs overlaid onto the ANT or ACC tSNE pseudo-time trajectory, respectively (units log_2_ CPM +1). **j**, tSNE visualization of the V1 visual cortex neurons pseudo-time trajectory colored by time point. **k**, tSNE V1 trajectory colored by pseudo-time. **l**, Kernel density estimate tSNE plots of V1 neurons across early-training, remote retrieval HR and remote retrieval LR.

**Extended data Fig. 7:**
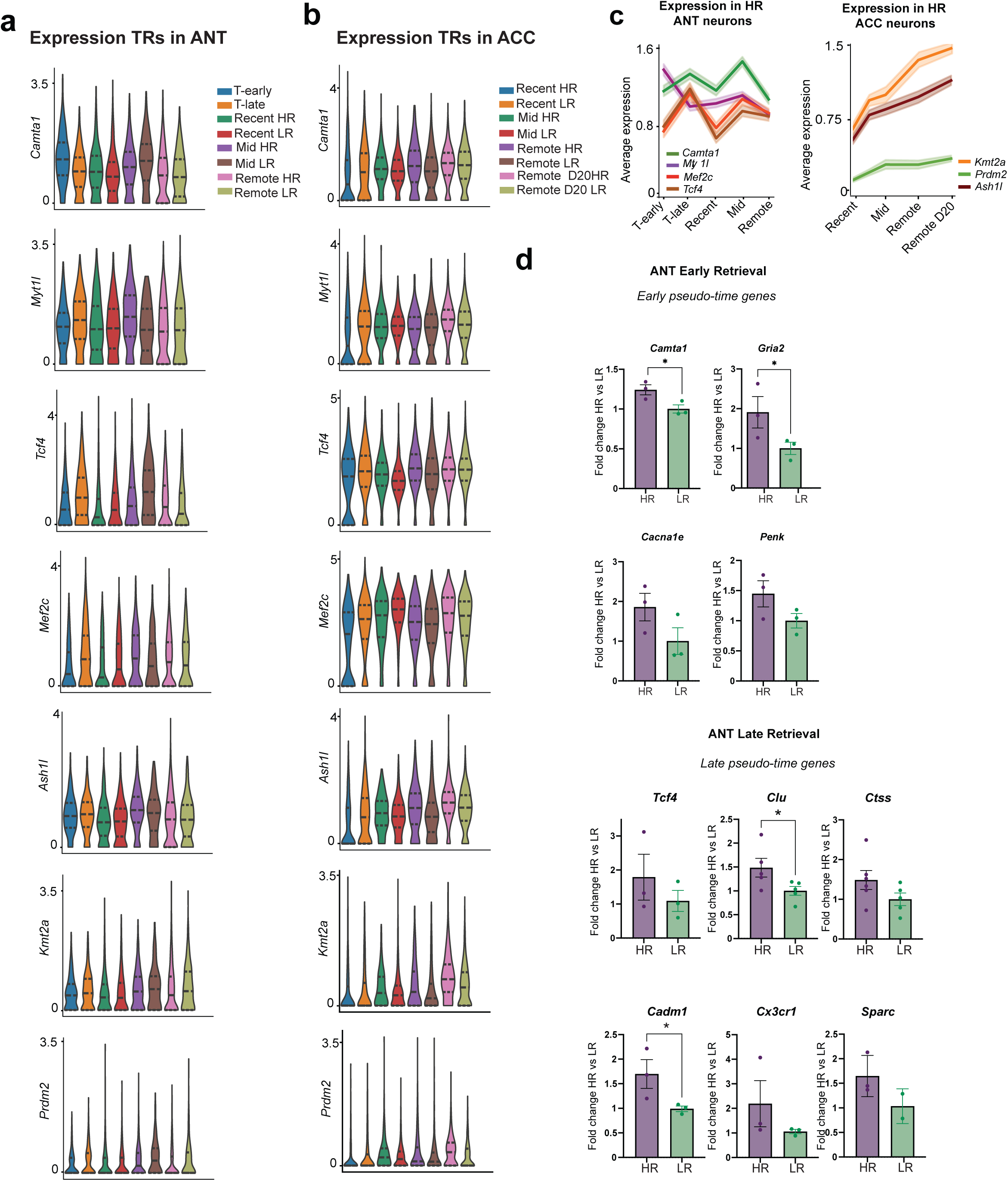
Expression of transcriptional regulators’ targets. **a**, Average expression of all transcriptional regulators in ANT neurons across conditions and time points (units log_2_ CPM +1). **b**, Average expression of all transcriptional regulators in ACC neurons across conditions and time points (units log_2_ CPM +1). **c**, Line plots of average expression of ANT (left) and ACC (right) transcriptional regulators in HR neurons only across behavioral time points (units log_2_ CPM +1). **d**, Relative expression of pseudo-time signature genes in ANT at early (top) and remote retrieval (bottom) between HR and LR samples, normalized to *Map2*, *n* = 3-6 biological replicates, \**P =* 0.0211 for *Camta1*; \**P =* 0.0495 for *Gria2*; \**P =* 0.0286 for *Clu*, \**P =* 0.0384 for *Cadm1,* unpaired t-test, mean± SEM shown. **e**, Relative expression of late pseudo-time signature genes in ACC between HR and LR samples, normalized to *Map2*, *n* = 5-6 biological replicates, mean± SEM shown.

**Extended data Fig. 8:**
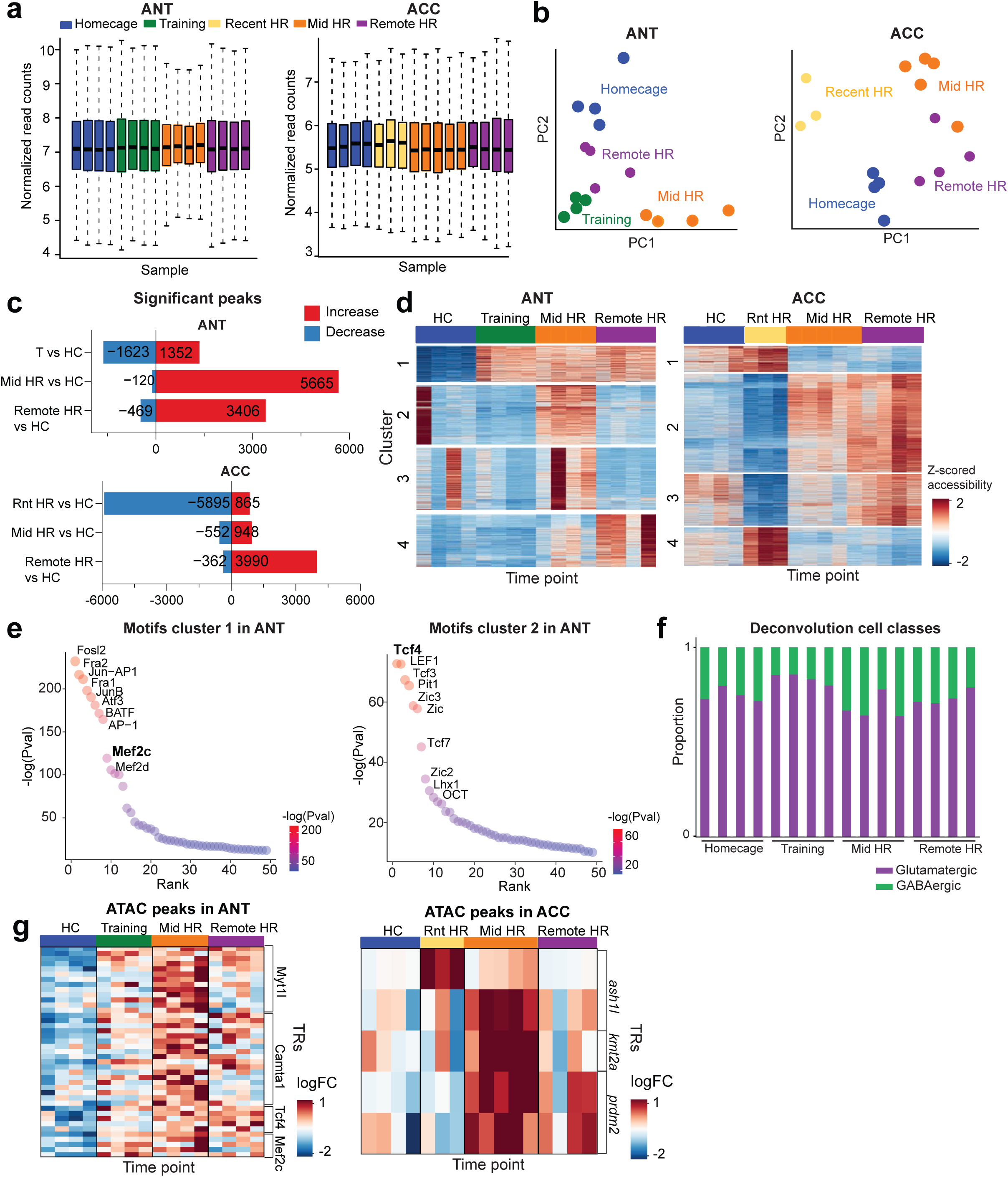
ATAC peaks of ACC transcriptional regulators’ modules remain accessible through remote retrieval. **a,** Boxplot of normalized read counts for all ANT (left) and ACC (right) ATAC-seq samples (*n* = 3-5 mice per time point). **b,** Principal component analysis (PCA) of samples from ANT (left) or ACC (right). Each point is the position of a single biological replicate (individual mouse), colored by the behavioral time point. **c,** Bar plot of significant peaks for HR conditions versus home-cage (HC) comparisons across time points in ANT (top) or ACC (bottom) neurons. **d,** Chromatin accessibility across ANT (left) or ACC (right) neurons. Rows are subsets of differentially accessible ATAC-seq peaks, and are organized into four clustered modules. Columns are ATAC-seq replicates and are colored by time point. **e,** Sorted rank plot of transcription factor motifs for cluster modules 1 and 2 from (**d**). for ANT neurons. **f**, Deconvolution analysis of ANT samples to estimate proportion of neuronal classes, assigned as vglut2 glutamatergic excitatory or GABAergic inhibitory. **g**, Z-scored accessibility of ATAC peaks of ANT (left) or ACC (right) TRs.

**Extended data Fig. 9:**
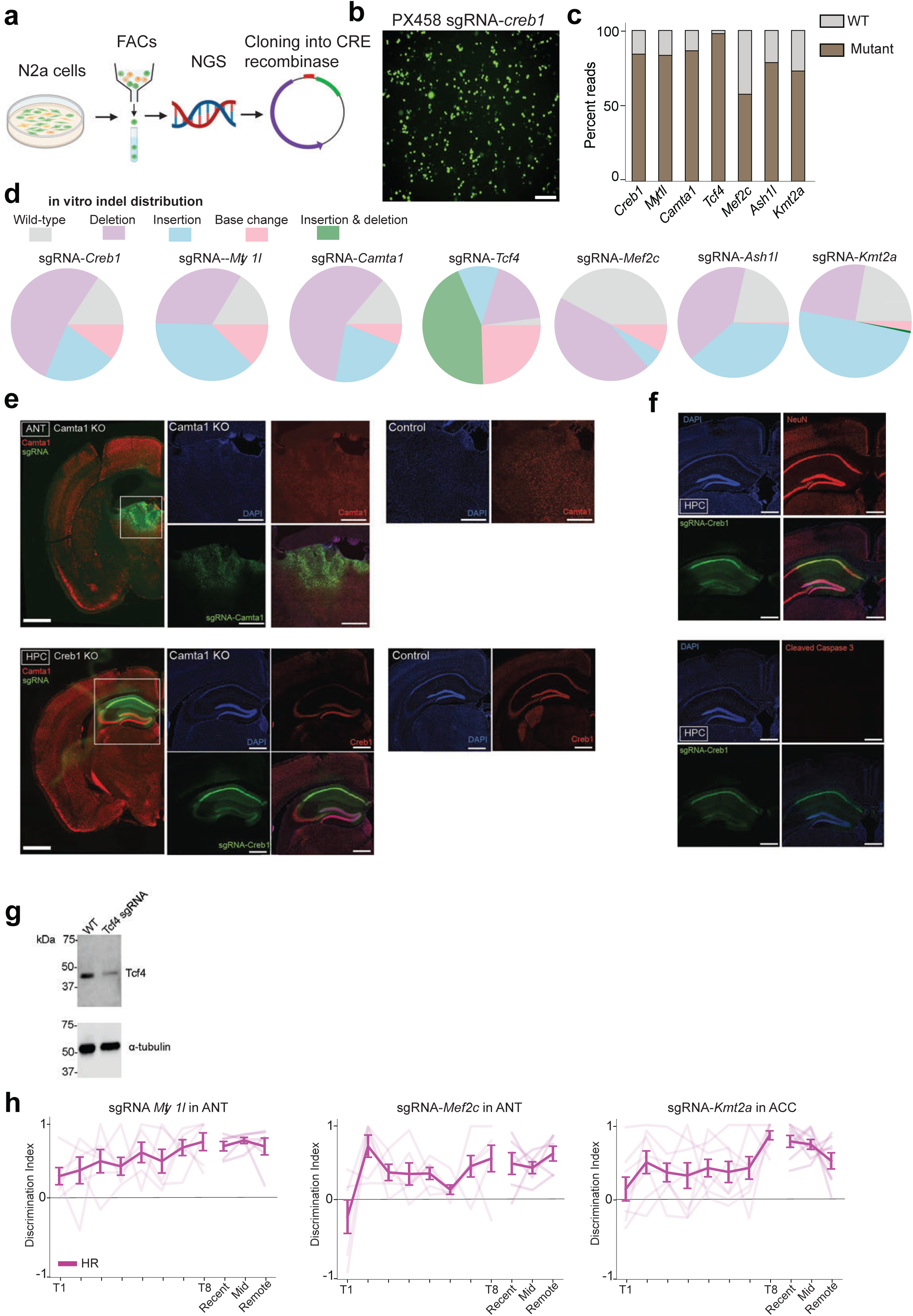
*In vitro* and *in vivo* sgRNA validation, and behavioral effects of TRs knockout. **a**, Schematic of *in vitro* CRISPR screen workflow. **b**, Neuro 2A cells transfected with sgRNA-spCas9-GFP, scale: 200um. **c**, Stacked bar plot of wildtype (WT) vs mutant percent reads of amplicon target regions from DNA collected from Neuro 2A cells post-transfection. **d**, Breakdown of Cas9-induced indel mutations in Neuro 2A cells expressing a sgRNA targeting one of the TRs. **e**, Top left: coronal slice from a *Camta1*-KO Rosa26-Cas9 mouse in ANT. Top middle inserts: 10X field of view from *Camta1*-KO displaying DAPI, CAMTA1 protein and sgRNA expression. Top right: 10X field of view from a control Rosa26-Cas9 mouse injected with CRE recombinase. Bottom left: coronal slice from a *Creb1*-KO in HPC. Bottom middle inserts: 10X field of view from *Creb1*-KO displaying DAPI, CREB1 protein and sgRNA expression. Bottom right: 10X field of view from a control Rosa26-Cas9 mouse injected with CRE recombinase. **f**, Immunofluorescent staining of NeuN or Cleaved Caspase 3 in Rosa26-Cas9 mouse injected with *Creb1*-sgRNA to evaluate extent of neuronal toxicity from AAV injection. Scale: 100µm in 4X, 50µm in 10X. **g**, Western blot validation of *Tcf4*-KO from ANT tissue compared to control animal. **h**, Discrimination indices for learning and recall performances in HR of *Rosa26LSL-spCas9-eGFP* mice expressing sgRNA targeting *Myt1l*, *Mef2c* or *Kmt2a* (*n* = 6-7 mice per cohort), individual data points shown (faded lines), with mean ± SEM (solid line). **g**, Example traces of F(z) from one control mouse on late training from ACC and ANT aligned to lick rate and task zones, three trial shown.

**Extended data Fig. 10:**
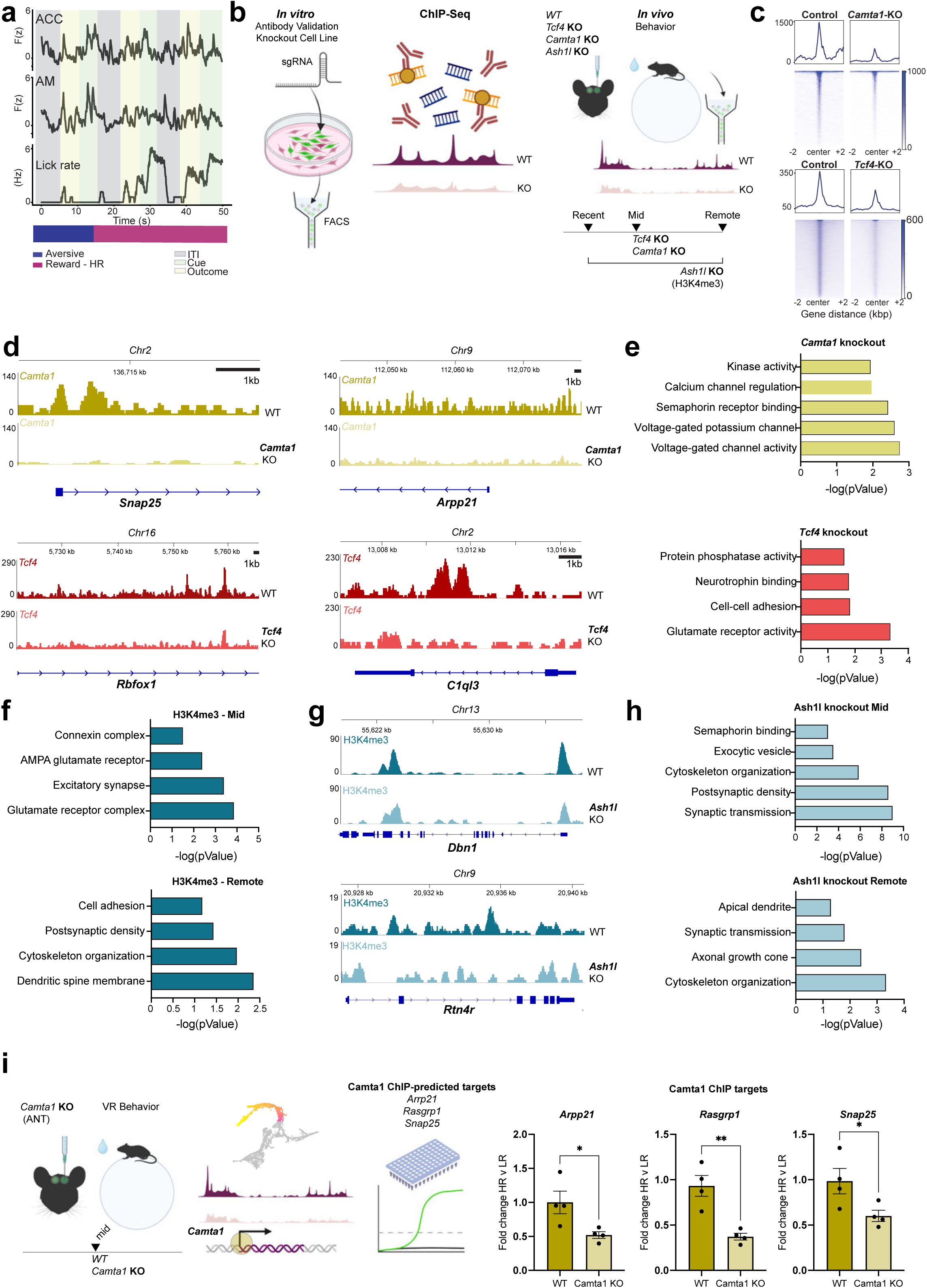
Mechanistic investigation of TRs by ChIP-seq and photometry. **a**, Example traces of F(z) from one control mouse from late training form ACC and ANT aligned to lick rate and task zones, three trials shown. **b**, Schematic of ChIP-Seq workflow: selection and validation of antibodies via generation of knockout N2A cell lines, followed by ChIP seq of animals collected from recent, mid, remote retrieval (for WT and *Ash1l* KO, H3K4me3 antibody) or from mid-retrieval (WT and *Camta1*/*Tcf4* KO). **c**, Tornado plots of overall ChIP-seq signal for controls and *Camta1*/*Tcf4* KO. **d**, Top: ChIP-seq signal at gene loci for *Camta1* (yellow) in WT vs. *Camta1*-KO mice at mid retrieval. Bottom: ChIP-seq signal at gene loci for *Tcf4* (red) in WT vs. *Tcf4*-KO mice at mid retrieval. Tracks show normalized read density (RPKM) and are color-coded by condition (darker shade for wildtype, lighter shade for KO). **e**, Gene ontology analysis derived from ChIP-seq differential peak analysis of WT vs. *Camta1*-KO (yellow) or WT vs. *Tcf4*-KO (red) at mid retrieval. **f**, Gene ontology analysis derived from H3K4me3 peaks (accessible chromatin) in wildtype animals at mid or remote retrieval. **g**, ChIP-seq signal at gene loci for H3K4me3 in WT or *Ash1l*-KO mice at remote retrieval. Tracks show normalized read density (RPKM) and are color-coded by condition (darker shade for wildtype, lighter shade for *Ash1l*-KO). **h,** Gene ontology analysis derived from H3K4me3 peaks from *Ash1l*-KO mice at mid and remote retrieval. **i**, Bar plots of downregulation of plasticity related gene targets in *Camta1*/-KO versus control animals (yellow), \**P* = 0.00159 for Arp21; \*\**P* = 0.0018 for Rasgrp1; \**P* = 0.0231 for Snap25, *n* = 4 biological replicates, errors bars are SEM.

